# The Utility of Ancestral and Derived Allele Sharing for Genome-Wide Inferences of Introgression

**DOI:** 10.1101/2022.12.02.518851

**Authors:** David Peede, Diego Ortega-Del Vecchyo, Emilia Huerta-Sánchez

**Author notes:** Corresponding Authors (DO-DV), (EHS).

## Abstract

The past decade has ushered in a resurgence of studies highlighting the importance of introgression throughout the Tree of Life. Several methods exist for detecting and quantifying introgression on a genomic scale, yet the majority of these methods primarily utilize signals of derived allele sharing between donor and recipient populations. In this study, we exploit the fact that introgression will not only result in derived allele sharing but also the reintroduction of ancestral alleles to derive new estimators of the admixture proportion. Using coalescent simulations, we assess the performance of our new methods and the methods proposed in *Lopez Fang et al. 2022* to assess the utility of incorporating shared ancestral variation into genome-wide inferences of introgression. Using coalescent theory, simulations, and applying our methods to human and canid data, we find that methods incorporating ancestral allele sharing are comparable to their derived allele sharing counterparts, in turn providing researchers with the opportunity to utilize more of the genomic signature of introgression.

## 1 Introduction

The field of evolutionary biology has seen a meteoric rise in the number of studies detecting and quantifying instances of introgression—the integration of foreign genomic blocks into a species’ genomic background—throughout the Eukaryotic Tree of Life (Dagilis et al., 2022) since the publishing of the draft Neanderthal genome in 2010 (Green et al., 2010). This is in large part due to the recent feasibility of next-generation sequencing technology, which has initiated a paradigm shift in how evolutionary biologists study introgression from comparing sequence similarity at some genetic markers between species to analyzing the whole genome via statistical methods—e.g., *Patterson’s D* —to make inferences of introgression by using site patterns as a proxy for genealogical relationships (Dagilis et al., 2022; Durand et al., 2011; Edelman & Mallet, 2021; Green et al., 2010). Recent years have ushered in an influx of studies demonstrating that introgression is not only more common than originally thought, but can also provide the raw genetic materials necessary for local adaptations (Hedrick, 2013; Huerta-Sánchez et al., 2014; Jagoda et al., 2022; Jones et al., 2018; Suarez-Gonzalez et al., 2018; Zhang et al., 2021), adaptive radiations (Malinsky et al., 2018; Meier et al., 2017; Pease et al., 2016; Richards & Martin, 2017; Stankowski & Streisfeld, 2015), and speciation (Bierne et al., 2013; Nelson et al., 2021; Rheindt & Edwards, 2011). However, even with keystone studies demonstrating that the evolutionary outcomes of introgression are diverse, it is challenging to assess its significance in evolution because the frequency of introgression across the Tree of Life is still unclear (Dagilis et al., 2022; Edelman & Mallet, 2021).

The most popular statistical methods for making genome-wide inferences about introgression exploit asymmetries in observed site patterns (Dagilis et al., 2022). Statistical tests for site pattern asymmetries count the number of segregating sites that have a configuration of shared derived and ancestral alleles which are informative of introgression in a set of taxa. One example of these tests is *Patterson’s D* (Equation 1), where two configurations of derived allele sharing site patterns are used to test for the presence of introgression (Durand et al., 2011; Green et al., 2010). Genome-wide tests of introgression based on observed site patterns are relatively straight-forward to implement, computationally efficient to calculate, and provide a unified approach for detecting and quantifying introgression across the Tree of Life as they can be applied to both phylogenetic and population genetic data sets. Due to the utility of site pattern-based metrics of introgression, recent years have seen a rapid proliferation of site pattern-based statistics designed to detect and quantify genome-wide levels of introgression (Durand et al., 2011; Hamlin et al., 2020; Lopez Fang et al., 2022; Martin et al., 2014; Pease & Hahn, 2015; Pfeifer & Kapan, 2019). However, until recently existing methods have strictly relied on the idea of derived allele sharing—where the donor and recipient populations both harbor the same derived allele—which only captures a small portion of the total genomic signature produced by introgression (Lopez Fang et al., 2022). Notably, *Lopez Fang et al. 2022* recently proposed the *D+* statistic, which leverages both the sharing of derived and ancestral alleles between donor and recipient populations to detect genomic windows harboring introgressed segments in the genome.

In this study we aim to assess if ancestral allele sharing site patterns are informative about introgression on a broader genomic scale. We first propose novel metrics to quantify the proportion of introgression that utilize ancestral allele sharing site patterns. Using coalescent simulations we assessed the performance of our newly proposed metrics and the metrics to detect introgression proposed by *Lopez Fang et al. 2022*. Lastly, we assessed the performance in empirical data by applying all the introgression metrics to publicly available human and canid data sets (1000 Genomes Project Consortium et al., 2015; Freedman et al., 2014). We find that in both simulated and empirical data, all introgression metrics perform comparably, allowing researchers the opportunity to use multiple lines of evidence across more segregating sites throughout the genome to make inferences about introgression.

## 2 New Approaches

In practice, most site pattern-based metrics of introgression assume a four-taxon tree—Newick format: (((*P1, P2*), *P3*), *O*); site pattern format: *(P1’s allelic state, P2’s allelic state, P3’s allelic state, O’s allelic state)*—where *P1* and *P2* represent potential recipients of introgression from the donor population *P3*, and *O* represents an outgroup used to polarize ancestral states, in which *A* denotes the ancestral allele and *B* denotes the derived allele. Thus, the *ABBA* site pattern indicates that the derived allele is shared between *P2* and *P3*, while the *BABA* site pattern indicates that the derived allele is shared between *P1* and *P3. Patterson’s D* (Equation 1) specifically tests for patterns of derived allele sharing between donor and recipient populations and a significant excess of either aforementioned site pattern is suggestive of introgression (Durand et al., 2011; Green et al., 2010)

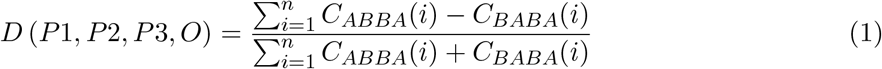

*C*_*ABBA*_(*i*) *− C*_*BABA*_(*i*)where *C*_*ABBA*_(*i*) and *C*_*BABA*_(*i*) are indicator variables that take a value of either zero or one conditioned on observing an *ABBA* or *BABA* site pattern at site *i* for all *n* sites where we have data for all four samples (Durand et al., 2011; Green et al., 2010). Due to its computational efficiency and ability to discriminate between coalescent histories of incomplete lineage sorting (ILS)—the process where two lineages fail to coalesce in their most recent common ancestral population—and introgression, *Patterson’s D* is the most widely used summary statistic to detect introgression (Dagilis et al., 2022). However, there is no theoretical or biological reason to believe that introgression will only result in the sharing of derived alleles. This is because introgressed genomic segments contain both new derived alleles and re-introduced ancestral alleles. Using this logic, *Lopez Fang et al. 2022* derived the expected branch lengths where a single mutation would generate a *BAAA* and *ABAA* site pattern respectively, which represents the ancestral allele sharing analogs to the *ABBA* and *BABA* site patterns. Notably, the authors demonstrated that the expected difference in the number of *ABBA* and *BABA* sites is equivalent to the expected difference in the number of *BAAA* and *ABAA* sites in the instantaneous unidirectional admixture (IUA) model of introgression—see the Appendix for the model description—providing the motivation to construct *D*_*anc*_ (Equation 2), a statistic that leverages ancestral allele sharing to detect local introgressed regions

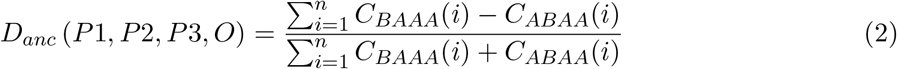

*C*_*BAAA*_(*i*) *− C*_*ABAA*_(*i*)where *C*_*BAAA*_(*i*) and *C*_*ABAA*_(*i*) are indicator variables that take a value of either zero or one conditioned on observing a *BAAA* or *ABAA* site pattern at site *i* for all *n* sites where we have data for all four samples. They also defined *D+*, a statistic (Equation 3) which leverages both the sharing of derived and ancestral alleles to strengthen inferences for detecting locally introgressed genomic regions

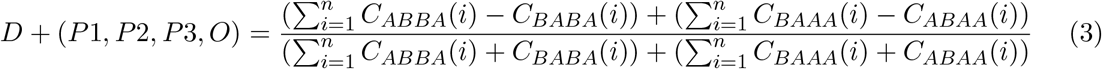

and found that *D+* was a more reliable statistic than *D* to detect introgression in small genomic windows (Lopez Fang et al., 2022). As researchers are also interested in quantifying the amount of introgression, here we construct estimators to infer the fraction of the genome shared through introgression—hereafter referred to as the admixture proportion. One approach to infer the admixture proportion is to assess the ratio of observed differences in allele sharing between the donor and recipient populations—i.e., the *P3* and *P2* populations respectively—in the numerator and the expected differences assuming the entire genome was introgressed—i.e., complete homogenization of allele sharing—from *P3* to *P2* in the denominator (Martin et al., 2014). This approach (Equation 4) was first described only in terms of derived allele sharing between recipient and donor populations in *Martin et al. 2015* as an alternative to the original estimator of the admixture proportion described in *Durand et al. 2011* where two pseudo-haploid genomes are created by randomly sub-sampling chromosomes from the *P3* individual. In the simplest case of only having four whole-genome alignments—which are monoploid—this pseudo-haploid approach is not possible, so *Martin et al. 2015* proposed to use the sampled *P3* sequence twice in the denominator to circumvent the need for creating pseudo-haploid genomes. The original estimator of the admixture proportion from *Martin et al. 2015* is *f*_*hom*_ (Equation 4)

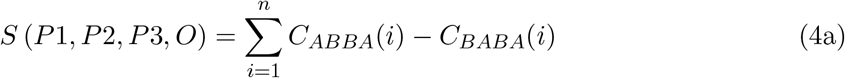

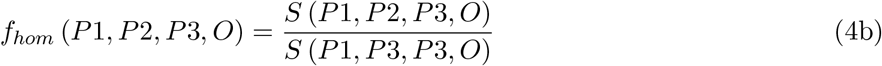

where homogenization is accomplished by replacing *P2* with *P3* in the assumed underlying species tree—i.e., in Newick format (((*P1, P3*), *P3*), *O*)—when computing the denominator. It should be noted that when written in terms of derived allele frequencies and in the most simplest case of only having four whole-genome sequences, *f*_*hom*_ is equivalent to *F4-ratio* (Patterson et al., 2012). Given that patterns of ancestral allele sharing between donor and recipient populations mirror those of their derived allele sharing counterparts we can re-derive *f*_*hom*_ in terms of *BAAA* and *ABAA* site patterns resulting in *f*_*anc*_ (Equation 5).

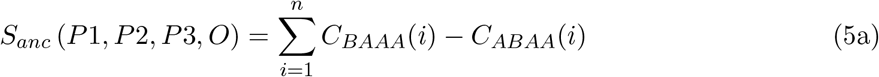

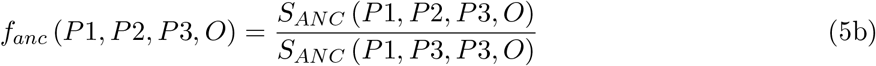

Lastly, given that introgressed segments are inherited in blocks consisting of both new derived alleles and re-introduced ancestral alleles one can formulate quantification metrics that consider all possible site patterns that represent allele sharing between recipient and donor populations; such as *f+* (Equation 6).

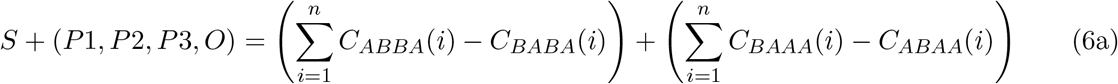

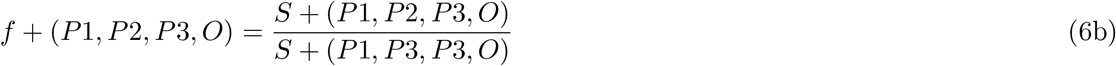

Note that all coalescent based expectations and equations written in terms of derived allele frequencies can be found in the Appendix.

## 3 Results

### 3.1 Ancestral & Derived Allele Sharing Site Pattern Differences Are Equivalent Across All Admixture Proportions

We first validated the theoretical result proposed by *Lopez Fang et al. 2022* that the (*ABBA – BABA*) and (*BAAA – ABAA*) site pattern differences are equivalent by simulating whole-genome sequences under an IUA model of introgression using parameters that reflect the demographic history of humans (Figure S1) using the coalescent simulator `msprime v1.1.1` (Baumdicker et al., 2022). For each simulation replicate, we sample one chromosome from each population, and computed the difference—i.e., (*ABBA – BABA*) – (*BAAA – ABAA*). We simulated 100 replicates under the demographic model to generate a distribution of the difference. We calculated the mean of the difference over the number of replicates and we take the absolute value of the mean—so that the mean difference is always positive. We then test to see if the mean is statistically significantly different from zero. For different values of the admixture proportion ranging from 0% to 50%, we find that the difference of site pattern differences does not significantly differ from zero under the IUA model (see Figure 1 and Table S4). This is also true in the case of a more complex demographic history—e.g. such as the one in *Ragsdale and Gravel 2019* —or when we consider a larger sample size (n=100), where the variance of the differences gets smaller (see Figure 1; Tables S4-S6 and S19-S20). These results suggest that the theoretical expectation is robust to sampling scheme, demographic history, and the amount of introgression, providing us with justification for constructing introgression metrics that incorporate ancestral allele sharing site patterns.

**Figure 1:**
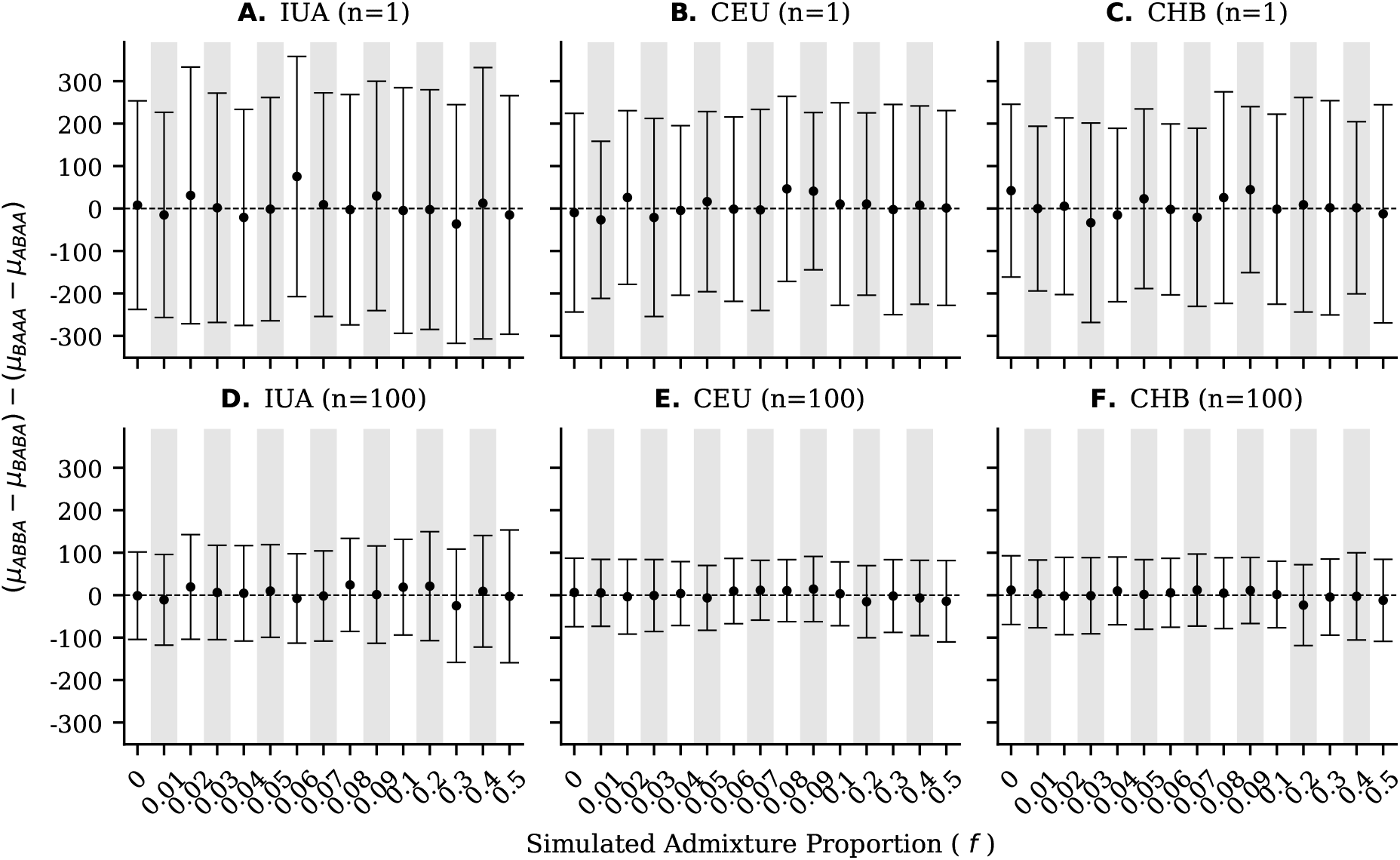
The mean (dots), standard deviation (error bars), and the theoretical expectation (dashed line and y = 0) for the difference of site pattern differences in simulations under an IUA model of introgression (panels **A** and **D**) and a more realistic model of human demography modified from *Ragsdale and Gravel 2019* (panels **B**-**C** and **E**-**F**) with a sampling scheme of n = 1 and n = 100 genomes from each of the potential recipient populations.

### 3.2 The Power To Detect Introgression Is Comparable Across All Metrics

Simulating under the IUA demographic model, we generated distributions of *D, D*_*anc*_, and *D+* and subsequently computed the mean simulated value per admixture proportion. Table S10 shows that the mean simulated values for these statistics match the values of *D, D*_*anc*_ and *D+* calculated using the theoretical expectations of the number of *ABBA, BABA, BAAA*, and *ABAA* site patterns (Equations A1-A3). We next assessed how the power to correctly detect introgression in simulated data differed between *D, D*_*anc*_, and *D+*. We define power as the number of replicates that significantly differ from zero out of 100 total simulated replicates per admixture proportion; to test for statistical significance we constructed 1000 bootstrapped genomes per simulation replicate and calculate each introgression dection metric, which were used to generate a *z* -distribution and subsequent *p-values*. Like *D, D+* and *D*_*anc*_ are able to correctly identify instances of introgression and we observe that the power to correctly identify instances of introgression is largely consistent across all metrics (Figure 2; Tables S7-S9). Notably, in simulations with no introgression (i.e., when *f = 0*) *D+* and *D*_*anc*_ either always incorrectly detect introgression at the same rate or lower rate as *D*, suggesting that on average *D+* and *D*_*anc*_ have a lower false positive rate (FPR) than *D*. Under the IUA model of introgression *D* and *D+* are always able to correctly identify introgression at comparable rates and outperform *D*_*anc*_. For more realistic demographic histories, on average *D* slightly outperforms *D+* for smaller amounts of introgression while both statistics tend to outperform *D*_*anc*_ (Tables S8-S9). Additionally, the power to correctly detect introgression always increased when the sample size increases in all models explored (Figure 2; Tables S7-S9).

**Figure 2:**
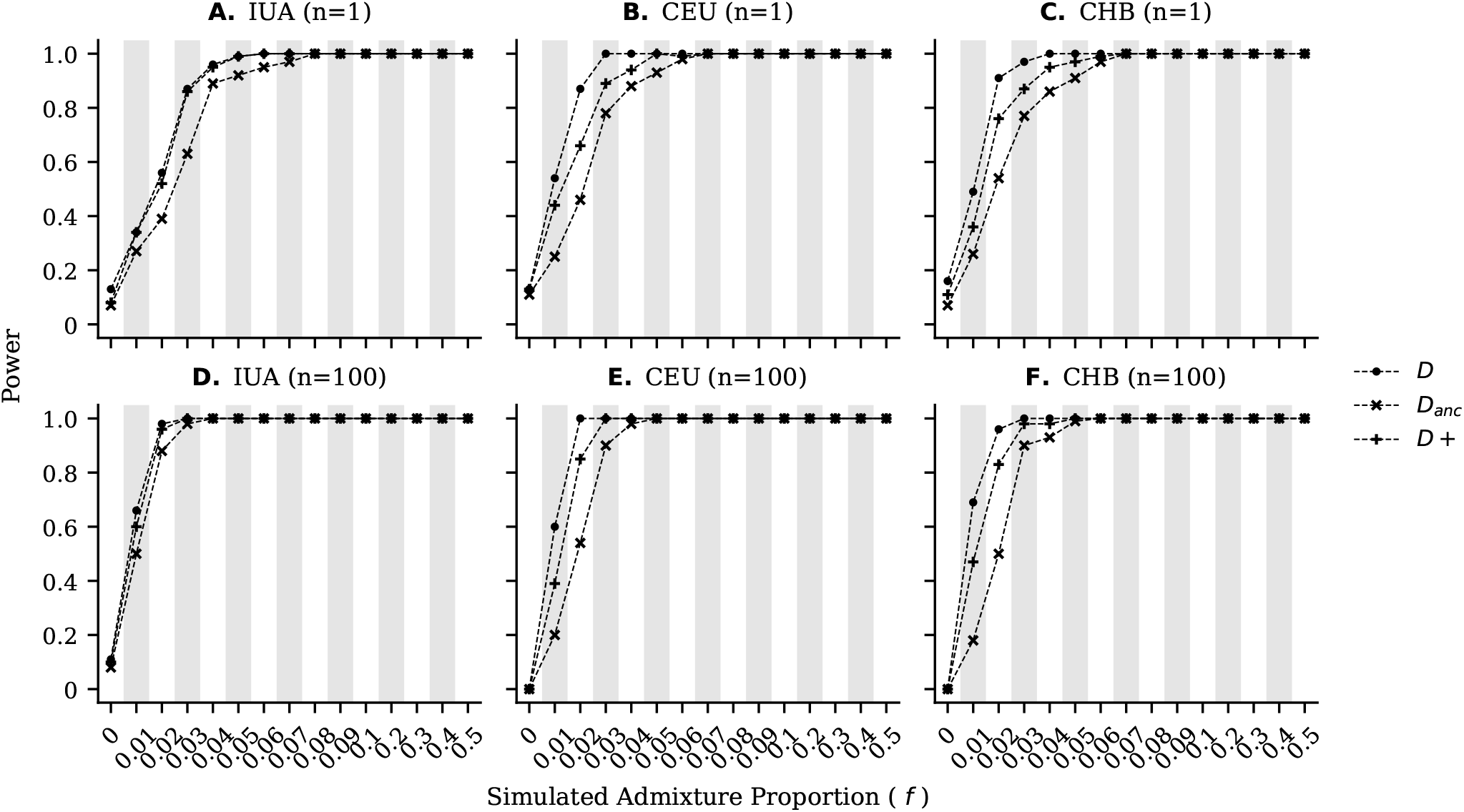
The power to correctly detect introgression in simulations under an IUA model of introgression (panels **A** and **D**) and a more realistic model of human demography modified from *Ragsdale and Gravel 2019* (panels **B**-**C** and **E**-**F**) with a sampling scheme of n = 1 and n = 100 genomes from each of the potential recipient populations. Note that the power at *f = 0* corresponds to the false positive rate since no introgression has occurred.

### 3.3 The Power To Quantify Introgression Is Nearly Identical Across All Metrics

Simulating under the IUA model with varying admixture proportions from 0% to 50% we generated distributions and computed the simulated mean values of *f*_*hom*_, *f*_*anc*_, and *f+*. Notably, the coalescent based expectations closely mirror the mean inferred admixture proportions across all simulations, providing evidence that *f*_*hom*_, *f*_*anc*_, and *f+* are all accurate metrics for quantifying introgression given the underlying IUA model (Figure 3; Table S16). We next evaluated the power of *f*_*hom*_, *f*_*anc*_, and *f+* to infer the true admixture proportion in simulated data under the assumption that a perfect estimator of the admixture proportion would infer the same value as the simulated admixture proportion—i.e., 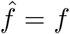. Under this assumption we calculated the root-mean-square-error (*RMSE*) and mean-absolute-error (*MAE*) of each quantification metric for all simulated admixture proportions. Despite that *f*_*hom*_, *f*_*anc*_, and *f+* all generally underestimate the true admixture proportion for our demographic models, the power to infer the admixture proportion is nearly identical across all metrics (Table S13-S15). In fact, Tables S16-S18 demonstrate that the mean inferred admixture proportions varied at most by 0.1% across all metrics for any given demographic model, sampling scheme, and simulated admixture proportion. However this is to be expected given that *f*_*hom*_, *f*_*anc*_, and *f+* all have the same coalescent based expectation—i.e., 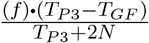 (see equationsA7-A9 in the Appendix).

**Figure 3:**
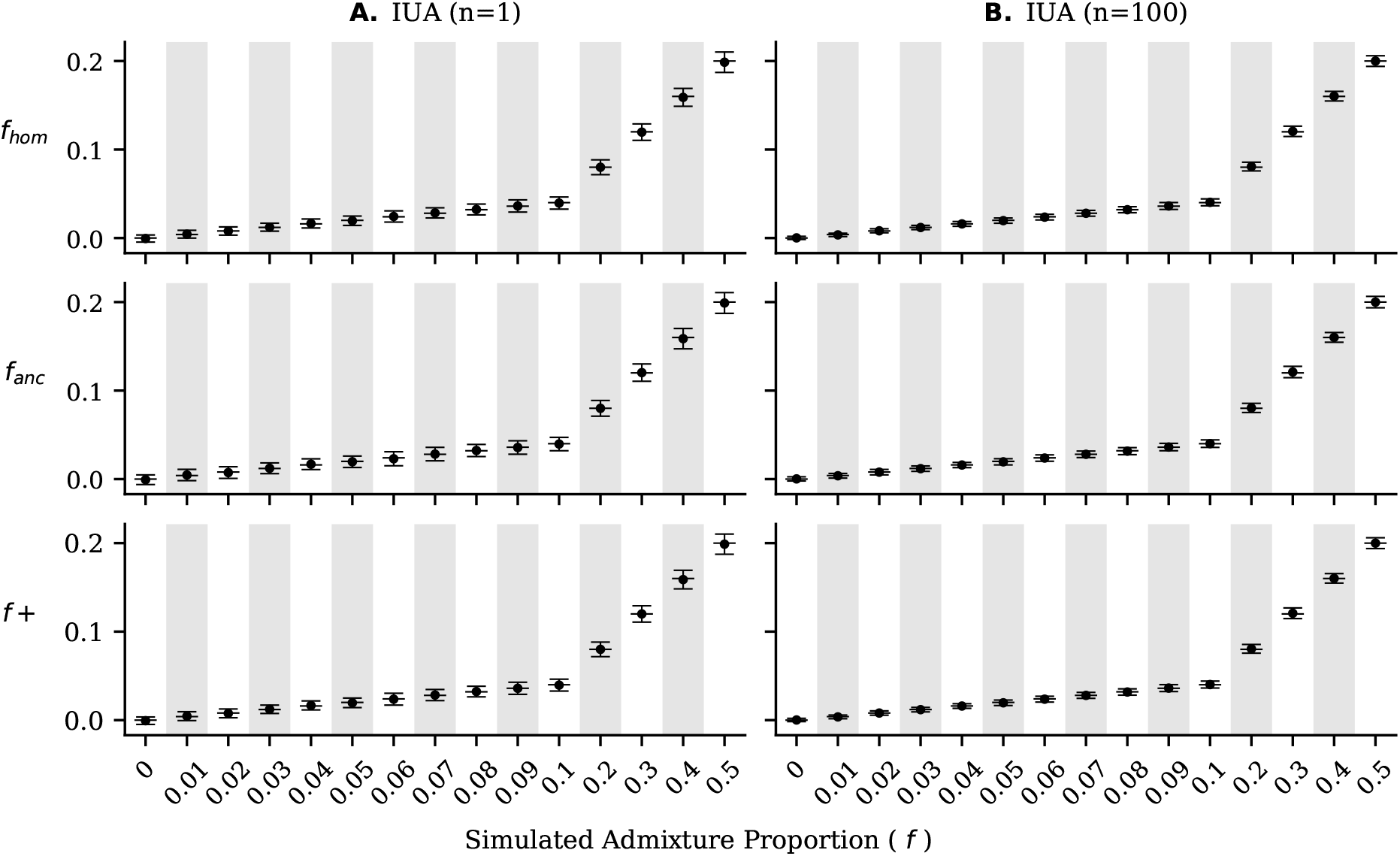
The mean (dots), standard deviation (error bars), and the theoretical expectation (solid horizontal lines) of *f*_*hom*_, *f*_*anc*_, and *f+* in simulations under an IUA model of introgression with a sampling scheme of n = 1 (panel **A**) and n= 100 (panel **B**) genomes from each of the potential recipient populations.

### 3.4 All Introgression Metrics Perform Similarly In Human & Canid Data

Before calculating *f*_*hom*_, *f*_*anc*_, and *f+* in empirical data, we wanted to investigate the behavior of ancestral and derived allele sharing site patterns. We first calculated *ABBA, BABA, BAAA*, and *ABAA* using the modern-day humans in the 1000 Genomes Project (TGP) and the high-coverage Altai Neanderthal genome (1000 Genomes Project Consortium et al., 2015; Prüfer et al., 2014). We first wanted to test if the difference of site pattern differences significantly deviated from zero by calculating site pattern counts for every non-African individual in the TGP and calculating the mean difference of site pattern differences per chromosome for every population. Concordant with our theoretical and simulated results, on average the mean difference of site pattern differences per chromosome was within two standard deviations of the theoretical expectation of zero (Figure 4). However, it should be noted that not all chromosomes were within two standard deviations, particularly, this was the case for chromosome 19 in every non-African population (Figure 4).

**Figure 4:**
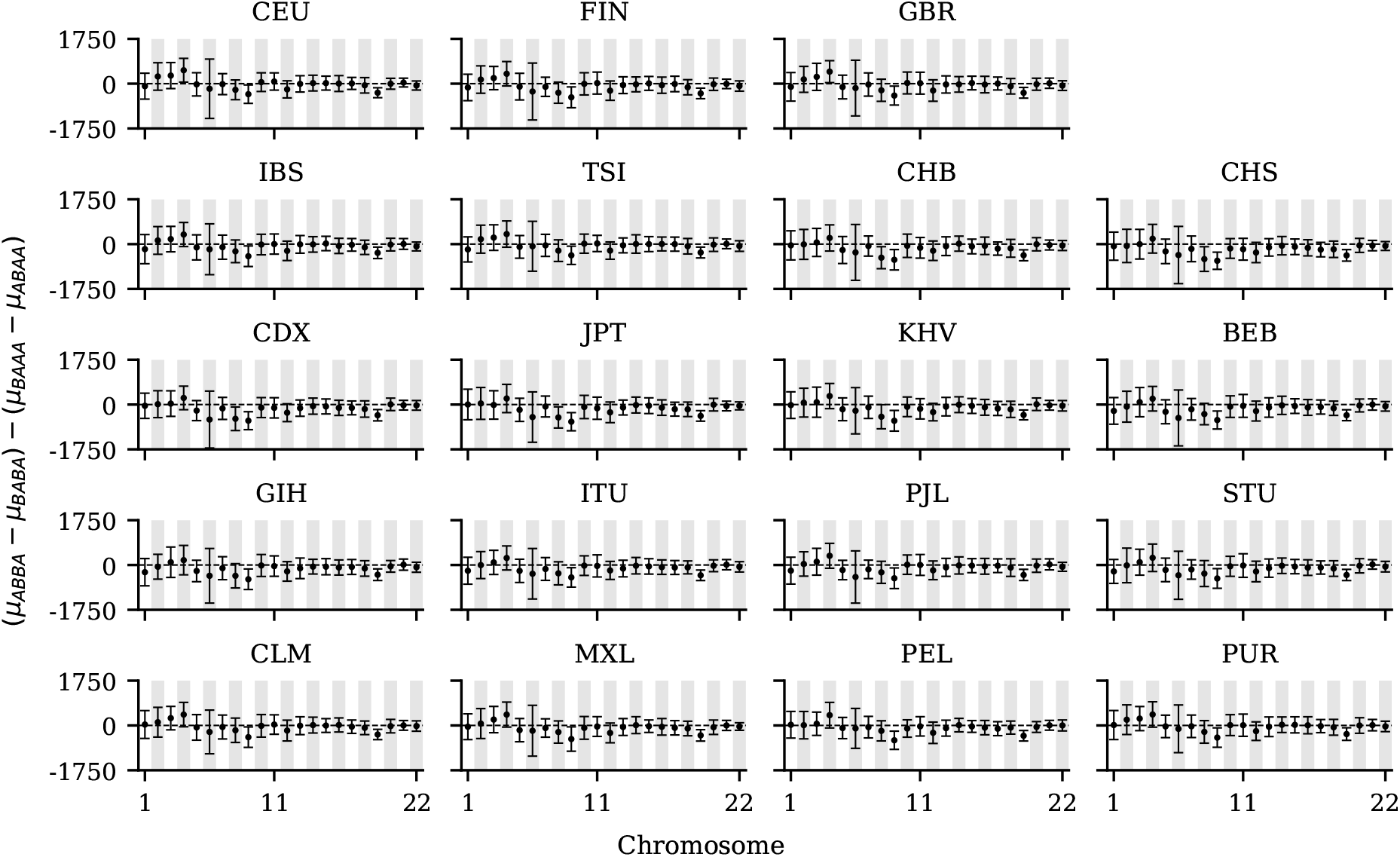
The mean (dots), two standard deviation (error bars), and the theoretical expectation (dashed line and y = 0) for the mean difference of site pattern differences among trios per chromosome for every non-African population in the 1000 Genomes Project.

Additionally, we computed the distribution of the difference across the whole genome (Figure 5) to assess if the genomic mean difference of site pattern differences significantly differed from zero in each population. After correcting for multiple comparisons, we found that only the CDX population did not meet our theoretical expectation (Figure 5; Table S21). Next, we inferred the Altai Neanderthal admixture proportion in each non-African individual and again concordant with our theoretical and simulated results we find that all estimates of the admixture proportion are concordant among trios with the mean *f*_*hom*_, *f*_*anc*_, and *f+* values all being within one standard deviation of one another within each non-African population (Figure 6; Table S22). As all the statistics studied here can also be defined as functions of allele frequency (see Equations A4-A6 and A10-A12), we also used the derived allele population frequencies to infer the presence and amount of introgression in every non-African population. Remarkably, we find that all *D, D*_*anc*_, and *D+* estimates are statistically significant after correcting for multiple comparisons and that estimates of *f*_*hom*_, *f*_*anc*_, and *f+* are nearly identical with nearly identical standard deviations (Table S23). Furthermore, estimates of the admixture proportion are in agreement with previous estimates of roughly 1% of Neanderthal introgression (Green et al., 2010; Mafessoni et al., 2020; Prüfer et al., 2017; Prüfer et al., 2014).

**Figure 5:**
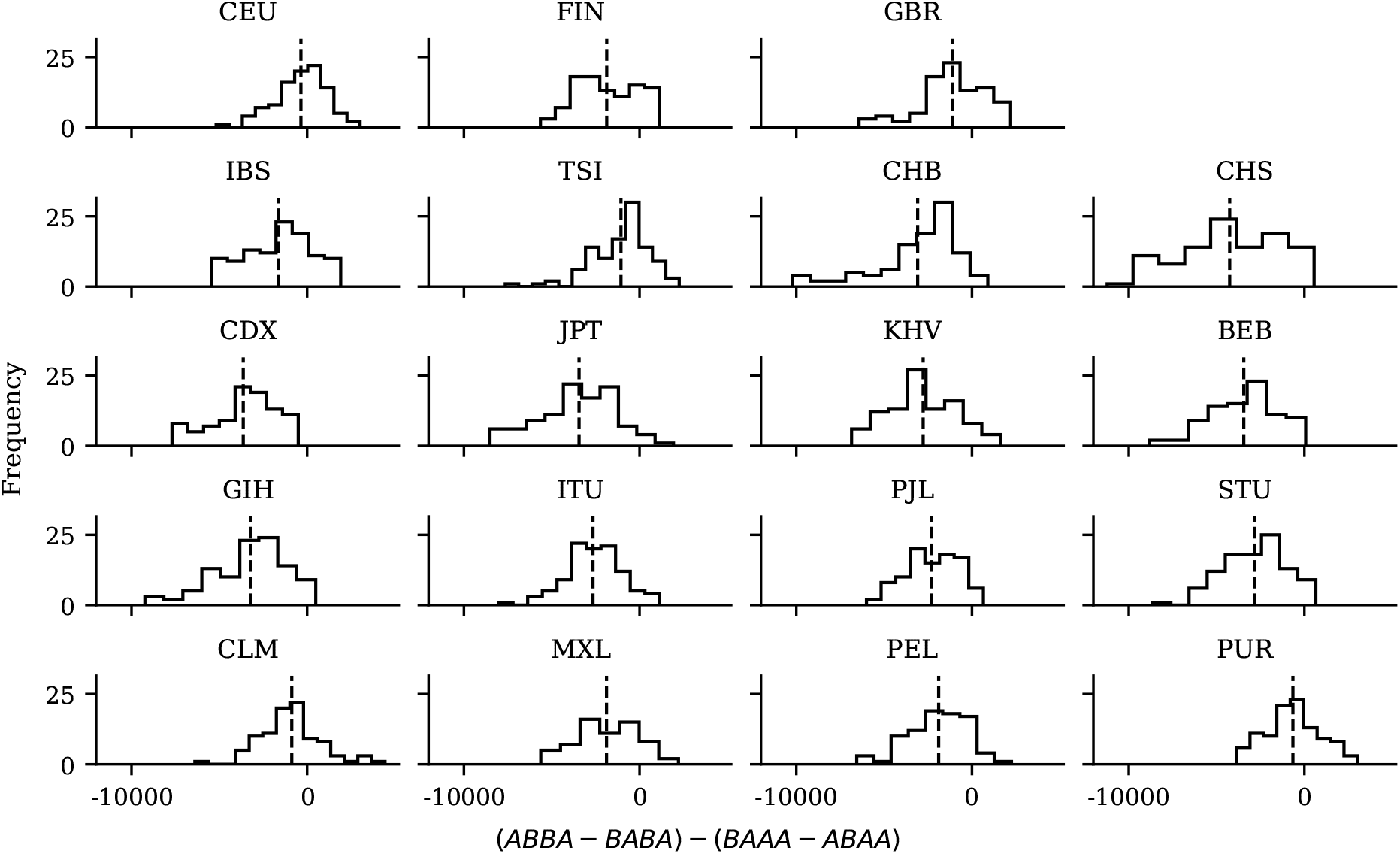
Distributions of difference of site pattern differences among trios for every non-African population in the 1000 Genomes Project. The black dashed line represents the genome-wide mean difference of site pattern differences among trios per non-African population.

**Figure 6:**
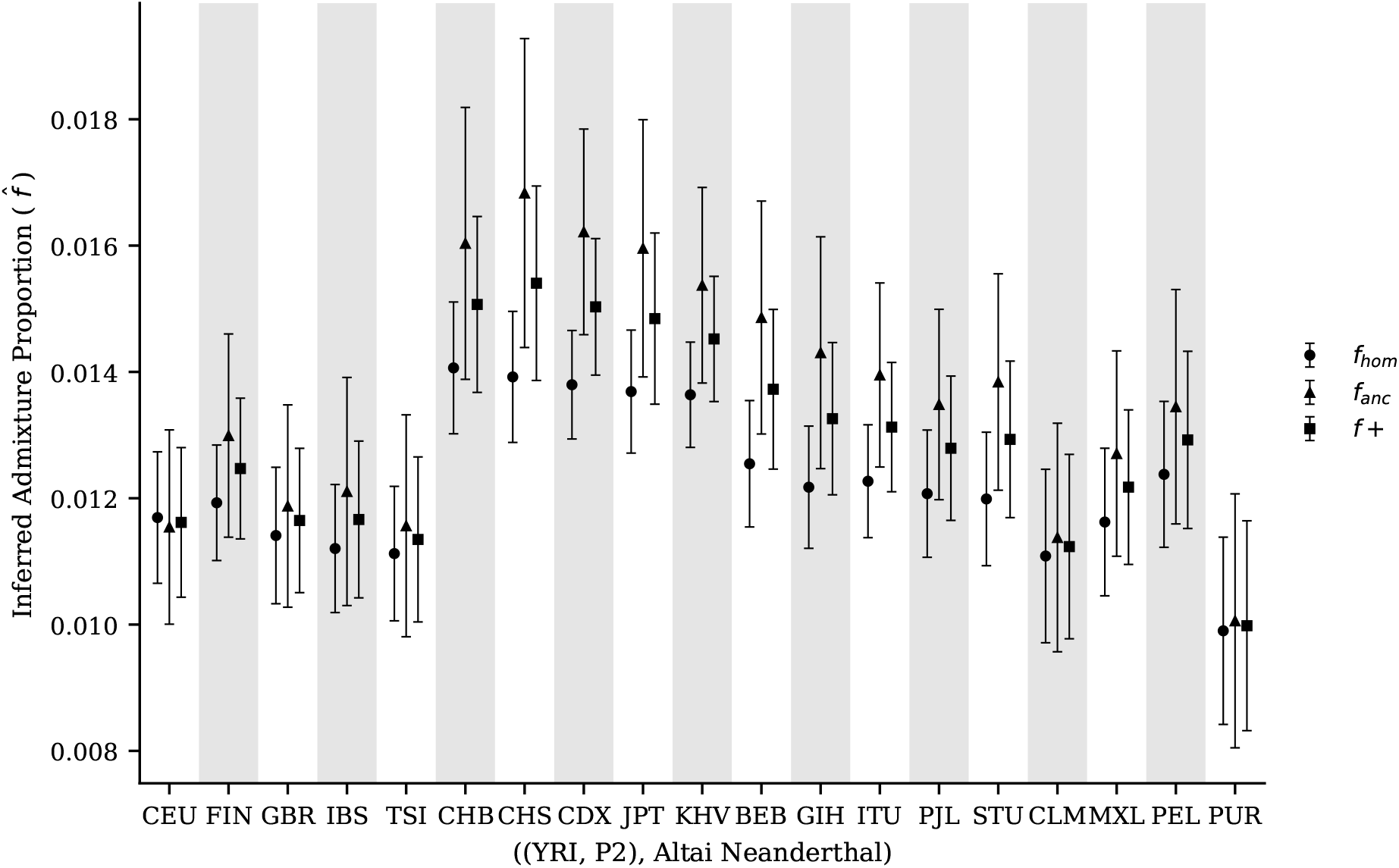
The mean (dots, triangles, and squares) and standard deviation (error bars) of *f*_*hom*_, *f*_*anc*_, and *f+* among trios for every non-African population in the 1000 Genomes Project.

We also sought out to apply our methods in a non-human data set, and we chose canid data to investigate the behavior of ancestral and derived allele sharing site pattern-based introgression metrics when only one individual per population is sequenced. To do so, we calculated all site patterns and introgression metrics from high-coverage whole-genome sequences for the ((Dingo, Basenji), Israeli Wolf) trio where we used the Golden Jackal as an outgroup—we chose this trio as it has the largest significant Z-score for *Patterson’s D* reported in the original paper (Freedman et al., 2014). It should be noted that it is highly unlikely that any biological system has experienced a demographic history that mirrors the IUA model which the underlying expectations for site pattern differences are based on; to this extent we propose a definition of *approximately equivalent*, which is defined as site pattern differences having the same sign and same order of magnitude. Notably, even though the demographic history of canids is known to violate the underlying IUA model, the site pattern differences are concordant with our definition of *approximately equivalent* (Table S24). Additionally, concordant with the original findings from *Freedman et al. 2014*, we find that all introgression detection metrics are statistically significant and that all inferred admixture proportions varied at most by 1.4% Israeli Wolf ancestry between quantification metrics (Table S24).

## 4 Discussion

The genomic era continues to provide evidence that post-divergence gene flow events are much more common than previously appreciated. Characterizing what are the effects and consequences of introgression across the tree of life requires providing robust tools that are able to efficiently and confidently detect and quantify such events. Here, in addition to using derived allele sharing between a putative donor and recipient population, we proposed to also incorporate ancestral allele sharing to construct estimators of the admixture proportion—e.g. *f*_*anc*_ and *f+*. In our study we benchmark *D, D*_*anc*_, *D+, f*_*hom*_, *f*_*anc*_, and *f+* which can be used to detect and quantify introgression. In particular, we use coalescent theory, simulations, and publicly available empirical data, to show that genome-wide site pattern-based tests of introgression that rely on ancestral, derived, and both patterns of allele sharing perform comparably, providing researchers the opportunity to corroborate claims of introgression from multiple metrics that exploit different genomic signals.

Our results show that *D, D*_*anc*_, and *D+* can detect the presence of introgression. On average *D*_*anc*_ has less power to detect introgression when compared to *D* and *D+*, but *D*_*anc*_ on average also exhibits the lowest FPR (Tables S7-S9). *D* tends to slightly have more power to detect introgression than *D+* but also tends to have higher FPR compared to *D+* (Tables S7-S9). In empirical data we found that *D, D*_*anc*_, and *D+* gave concordant results, suggesting that all of the aforementioned statistics are appropriate for detecting introgression on genome-wide data sets. The power to correctly infer the admixture proportion suffers from theoretical constraints. Specifically, *f*_*hom*_, *f*_*anc*_, and *f+* all have the same theoretical expectation of 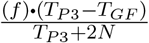 (see equations A7-A9 in the Appendix), which makes it clear that we will always underestimate the admixture proportion. In simulated data the mean *f*_*hom*_, *f*_*anc*_, and *f+* values closely mirrored the theoretical expectations (Figure 3; Table S13) and all quantification metrics exhibit nearly identical *RMSE* and *MAE* with respect to the simulated admixture proportion (Tables S13-S15). Additionally, in human data not only do we observe nearly identical estimates of the admixture proportions between all metrics studied here, but also nearly identical standard deviations for both trio and population level estimates (Figure 3; Table S23 and S22). Taken together we propose that researchers make use of both derived and ancestral allele sharing site pattern-based tests of introgression to make more thorough assessments about the presence and amount of introgression. To this extent we also developed a lightweight `python` package `RIPTA` (**R**e-evaluating **I**ntrogression site **P**atterns **T**hrough **A**ncestral alleles) to calculate all site patterns and introgression metrics studied in here.

Our theoretical and simulation results have shown that the expectation of site pattern differences— i.e., (*ABBA – BABA*) – (*BAAA – ABAA*)—is zero. This is true even when we consider complex demographic histories (Figure 1). When we count these site patterns using human and archaic genomes, we found that most non-African chromosomes in the TGP do not depart from this theoretical expectation (Figures 4-5; Table S21). One exception is chromosome 19 (see Figure 4), where the difference of (*BAAA – ABAA*) is larger than (*ABBA – BABA*), and this is true across all non-African populations. We tested to see if more complex demographic histories that have been proposed for human populations could lead to deviations from zero. For example, we tried incorporating population expansions and contractions, multiple pulses of introgression, and a dilution in the recipient population, and none of these models result in a difference that deviates from zero (Tables S5-S6 and S19-S20).

Another explanation for this observation, could be a component of evolution that our simulations ignored, such as natural selection. Studies have shown that Neanderthal alleles were originally costly on the modern human genetic background due to genetic load (Harris & Nielsen, 2016; Kim et al., 2018). It has also been shown that the effects of Neanderthal ancestry is negatively correlated with exon density under an additive fitness model (Kim et al., 2018). Given that chromosome 19 has the highest gene density in the genome (Grimwood et al., 2004), we hypothesize that the effects of natural selection may influence these site patterns. However, while some studies have shown that derived archaic alleles were likely mostly deleterious in modern humans, we know very little about what evolutionary forces (if any) acted on ancestral alleles that were re-introduced into modern humans with archaic introgression. We acknowledge that the *BAAA* and *ABAA* site patterns are likely to be more sensitive to differences in mutations rate in the two sister populations—i.e., *P1* and *P2* —than *ABBA* and *BABA* sites. This might be contributing to observation that the difference of (*BAAA – ABAA*) is larger than (*ABBA – BABA*), where more *BAAA* sites could indicate a higher mutations rate in *P1* vs *P2*. As this is only observed in chromosome 19, we think it is more likely that other evolutionary forces might be contributing to this signature, but more work is needed to characterize what can lead to deviations from zero.

In summary, we have provided further evidence that ancestral allele sharing site patterns are informative for detecting introgression on a genome-wide scale and are robust to realistic models of human demography. Additionally, we have derived novel estimators of the admixture proportion and demonstrated that these methods provide reliable estimates of the admixture proportion. Applying it to empirical data in humans suggests that chromosome 19 is an outlier with respect to the difference in ancestral allele sharing site patterns. None of the demographic histories that we simulated lead to deviations, so we hypothesize that other non-neutral processes may be at play. More work to investigate how natural selection acted on ancestral alleles that were reintroduced into human populations through introgression with archaic humans might provide insights into how evolutionary forces have shaped introgressed genetic variation.

## 5 Materials and Methods

All annotated code to recreate all analyses and annotated Jupyter Notebooks with walk-throughs of how to perform all analyses described in this paper can be found at https://github.com/David-Peede/anc-der-intro-proj. The RIPTA package to calculate site patterns and all introgression metrics studied here can be found at https://github.com/David-Peede/RIPTA.

### 5.1 Simulation Design

All demographic models were quality controlled using msprime v1.1.1 and visualized using demesdraw v0.3.0 (Baumdicker et al., 2022; Gower et al., 2022). All demes .yaml files with demographic histories needed to reproduce our results can be found https://github.com/David-Peede/anc-der-intro-proj/tree/main/simulations/yamls.

#### 5.1.1 IUA Model

To assess the statistical power of all introgression metrics discussed in this study we simulated whole-genome sequences based on a three-taxon tree following IUA model described in *Durand et al*., *2011*, and in *Green et al*., *2010*. Using the coalescent simulator `msprime v1.1.1` we simulated 100 Mb genomes under a demographic model with the following parameters: *T*_*P3*_ = 16,000 generations ago, *T*_*P2*_ = 4,000 generations ago, *T*_*GF*_ = 1,600 generations ago, constant and equal *N*_*e*_ = 10,000 for all populations, *r* = 1*e*^-8^ per bp per generation, and *µ* = 1.5*e*^-8^ per bp per generation under a Jukes-Cantor substitution model (Baumdicker et al., 2022). Our parameter space was constructed to reflect a simplified version of human evolution with introgression from the Neanderthal population into the Eurasian population as was originally described in *Racimo et al*., *2017*. The divergence times, timing and direction of introgression, and mutation and recombination rates are all direct reflections of the original parameter space, and represent plausible parameter values for the well studied case of Neanderthal introgression (Racimo et al., 2016). The only model parameter varied was the simulated admixture proportion *f ∈ {*0, 0.01, 0.02, 0.03, 0.04, 0.05, 0.06, 0.07, 0.08, 0.09, 0.1, 0.2, 0.3, 0.4, 0.5*}* which was used to determine the behavior of the introgression metrics studied here, for both plausible and theoretical amounts of introgression. For each of the 15 admixture proportions, 100 replicate simulations were performed, either n = 1 or n = 100 simulated genomes were sampled per potential recipient population per replicate, and one simulated genome was sampled from the donor population, which were then used to calculate site pattern counts and all introgression metrics described in this paper.

#### 5.1.2 Non-IUA Models

To determine the statistical power of all introgression metrics in a more realistic model of human demographic history, we used the used the demographic model proposed by *Ragsdale and Gravel 2019* implemented in `stdpopsim v0.1.2` (Adrion et al., 2020; Ragsdale & Gravel, 2019). To make this model more comparable to our analyses using the IUA model we modified the model (see Figure S2) to consist of discrete pulses of introgression—one pulse into the Eurasian population, and one pulse into CEU and CHB populations respectively—by taking the midpoint of the two periods of continuous bidirectional gene flow described in the original model. The simulation design for the modified *Ragsdale and Gravel 2019* model is the exact same as described in the previous section. Additionally, we ran all sets of analyses on the original model (see Figure S3) using the original estimates of bidirectional gene flow (Ragsdale & Gravel, 2019). Lastly, to assess if the difference of site pattern differences would significantly deviate from zero we chose to simulate multiple pulses of introgression, ghost introgression, and a dilution event from a so called “Basal Eurasian” population (see Figures S4-S5). We used demographic parameters described in *Rogers and Bohlender 2015, Villanea and Schraiber 2019*, and *Ragsdale and Gravel 2019* to run 100 replicate simulations per all pairwise possibilities of plausible Neanderthal *f*_*NEA*_ *∈ {*0, 0.005, 0.01, 0.015, 0.02*}* and Denisovan *f*_*DEN*_ *∈ {*0, 0.005, 0.01, 0.015, 0.02*}* admixture proportions for sampling schemes of n = 1 and n = 100 simulated genomes from the potential recipient populations, and one simulated genome from the Neanderthal population (Ragsdale & Gravel, 2019; Rogers & Bohlender, 2015; Villanea & Schraiber, 2019).

### 5.2 Assessing Introgression Metrics In Simulated Data

For all 100 replicate simulations per admixture proportion we assessed the theoretical expectation that (*ABBA – BABA*) is equivalent to (*BAAA – ABAA*) by computing the difference of site pattern differences—i.e., (*ABBA – BABA*) – (*BAAA – ABAA*)–per replicate to construct a *z* -distribution. We then assessed if the absolute value of the mean difference of site patterns differences significantly differed from the theoretical expectation of zero using the `scipy.stats.sf` function in `scipy v1.7.2`, and corrected for multiple comparisons using the Bonferroni correction (Virtanen et al., 2020). To determine the significance of introgression detection metrics—i.e., *D, D*_*anc*_, and *D+*—for each simulation replicate we built a bootstrapped distribution of 1,000 bootstrapped replicates, where each bootstrapped replicate consists of concatenating 1,000 100 kb windows that were randomly sampled with replacement to build 100 Mb bootstrapped genomes for each focal population. The bootstrapped genomes were then used to construct *z* -distributions of *D, D*_*anc*_, and *D+* per replicate, and a nominal *p-value* of *<* 0.05 was used as the significance threshold. The power to detect introgression using *D, D*_*anc*_, and *D+* was then determined by the proportion of replicates that were statistically significant per admixture proportion. The power to accurately infer the true—i.e., simulated—admixture proportion using *f*_*hom*_, *f*_*anc*_, and *f+* was assessed by calculating root-mean-square-error (*RMSE*) and mean-absolute-error (*MAE*) using the following equations:

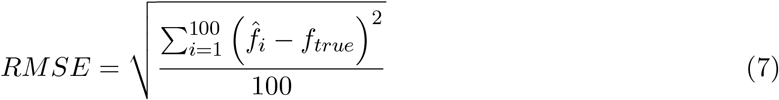

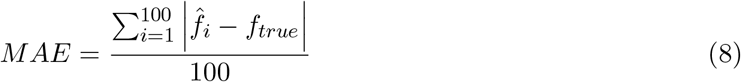

where 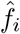 represents the inferred admixture proportion for the *i*^*th*^ replicate simulation out of 100 total replicates and where *f*_*true*_ represents the admixture proportion used in the simulation to infer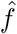. We chose to assess the performance of each introgression quantification metric using both the *RMSE* and *MAE* because both values are in the same units as the admixture proportion and because both quantify the amount of dispersion in the estimates of the inferred admixture proportion around the simulated admixture proportion where the *RMSE* values gives more weight to inferred estimates that have the large amounts of error while *MAE* values equally weight the amount of error in all inferred estimates.

### 5.3 Human Application

The all sites VCF files for the high-coverage genome of the Altai Neanderthal were downloaded from https://www.eva.mpg.de/genetics/genome-projects, the curated and imputed high quality genotypes from the phase three release of the 1000 Genomes Project (TGP) in VCF format was download from http://ftp.1000genomes.ebi.ac.uk/vol1/ftp/release/20130502, and the ancestral allele calls in fasta format for the Hg19 assembly using the Enredo, Pecan, Ortheus (EPO) pipeline was download from http://ftp.ensembl.org/pub/release-74/fasta/ancestral alleles (1000 Genomes Project Consortium et al., 2015; Herrero et al., 2016; Prüfer et al., 2014). The autosomal TGP and Altai Neanderthal VCF files were merged using the merge command from the bcftools v1.13 package and the resulting merged VCF files were then filtered to only contain sites with no missing data, an ancestral allele call, and bi-allelic SNPs with a mapping quality of 25 or higher and a genotype quality of 40 or higher (Li, 2011). Site patterns and all introgression metrics were then calculated for every non-African individual using the following configuration *P1* = NA18486 (a randomly chosen individual from Yoruba in Ibadan, Nigeria), *P2* = non-African individual, *P3* = Altai Neanderthal, and *O* = EPO ancestral allele call. Statistical tests based on the difference of site pattern difference distributions per non-African TGP population were carried out in the same manner as described in previous methods subsection for both individual chromosomes and the entire genome. Next, site patterns and all introgression metrics were additionally calculated from derived allele frequencies using the following configuration *P1* = Yoruba in Ibadan, Nigeria (YRI), *P2* = non-African population, *P3* = Altai Neanderthal, and *O* = EPO calls. Lastly, for the population level comparisons, the significance of introgression detection metrics—including correcting for multiple comparisons using the Bonferroni correction—and standard deviations for the introgression quantification metrics were determined from bootstrapped distributions consisting of 1,000 replicates, which were generated from 1,000 bootstrapped genomes per population, where a single bootstrapped genome was constructed from concatenating 303 randomly sampled with replacement 10 Mb windows.

### 5.4 Canid Application

The merged VCF files for the high-coverage canid data-set were downloaded from https://doi.org/10.5061/dryad.sk3p7 (Freedman et al., 2014). The autosomal VCF files were then filtered by the genomic-feature (`GF=1`) flag using the `view` command from the `bcftools` v1.13 package. Additionally, when performing calculations we only used sites where all focal individuals passed their respective sample filters—i.e., `SF=1` flag. We calculated site patterns and all introgression metrics for the configuration *P1* = Dingo, *P2* = Basenji, *P3* = Israeli Wolf, and *O* = Golden Jackal because the aforementioned trio had the highest degree of confidence for a significant *Patterson’s D* reported in the original paper (Freedman et al., 2014). Lastly, the significance of introgression detection metrics and standard deviations for the introgression quantification metrics were determined from bootstrapped distributions consisting of 1,000 replicates, which were generated from 1,000 bootstrapped genomes per individual canid, where a single bootstrapped genome was constructed from concatenating 220 randomly sampled with replacement 10 Mb windows.

## 6 Data Availability

All annotated code and previously published publicly available data sets needed to reproduce the entirety of this manuscript can be found at the url links described in the methods section.

## 8 Acknowledgements

The authors are grateful to Mia Miyagi, Andrius Dagilis, Mark Hibbins, Patrick McKenzie, Bastian Pfeifer, Durrell Kapan, Daniel Weinreich, and Yaniv Brandvain for many insightful discussions about coalescent theory and introgression. We would also like to thank Elizabeth Chevy, Emmanuel D’Agostino, Ria Vinod, Cole Williams, and the members of the Huerta-Sánchez laboratory for riveting discussions that helped improve this manuscript.

## References

1000 Genomes Project Consortium, Auton, A., Brooks, L. D., Durbin, R. M., Garrison, E. P., Kang, H. M., Korbel, J. O., Marchini, J. L., McCarthy, S., McVean, G. A., & Abecasis, G. R. (2015). A global reference for human genetic variation. Nature, 526 (7571), 68–74.

Adrion, J. R., Cole, C. B., Dukler, N., Galloway, J. G., Gladstein, A. L., Gower, G., Kyriazis, C. C., Ragsdale, A. P., Tsambos, G., Baumdicker, F., Carlson, J., Cartwright, R. A., Durvasula, A., Gronau, I., Kim, B. Y., McKenzie, P., Messer, P. W., Noskova, E., Ortega-Del Vecchyo, D., … Kern, A. D. (2020). A community-maintained standard library of population genetic models. Elife, 9.

Baumdicker, F., Bisschop, G., Goldstein, D., Gower, G., Ragsdale, A. P., Tsambos, G., Zhu, S., Eldon, B., Ellerman, E. C., Galloway, J. G., Gladstein, A. L., Gorjanc, G., Guo, B., Jeffery, B., Kretzschumar, W. W., Lohse, K., Matschiner, M., Nelson, D., Pope, N. S., … Kelleher, J. (2022). Efficient ancestry and mutation simulation with msprime 1.0. Genetics, 220 (3).

Bierne, Gagnaire, & David. (2013). The geography of introgression in a patchy environment and the thorn in the side of ecological speciation. Curr. Zool.

Dagilis, A. J., Peede, D., Coughlan, J. M., Jofre, G. I., et al. (2022). A need for standardized reporting of introgression: Insights from studies across eukaryotes. Evolution.

Durand, E. Y., Patterson, N., Reich, D., & Slatkin, M. (2011). Testing for ancient admixture between closely related populations. Mol. Biol. Evol., 28 (8), 2239–2252.

Edelman, N. B., & Mallet, J. (2021). Prevalence and adaptive impact of introgression. Annu. Rev. Genet., 55 (1), 265–283.

Freedman, A. H., Gronau, I., Schweizer, R. M., Ortega-Del Vecchyo, D., Han, E., Silva, P. M., Galaverni, M., Fan, Z., Marx, P., Lorente-Galdos, B., Beale, H., Ramirez, O., Hormozdiari, F., Alkan, C., Vilà, C., Squire, K., Geffen, E., Kusak, J., Boyko, A. R., … Novembre, J. (2014). Genome sequencing highlights the dynamic early history of dogs. PLoS Genet., 10 (1), e1004016.

Gower, G., Ragsdale, A. P., Bisschop, G., Gutenkunst, R. N., Hartfield, M., Noskova, E., Schiffels, S., Struck, T. J., Kelleher, J., & Thornton, K. R. (2022). Demes: A standard format for demographic models. Genetics, 222 (3).

Green, R. E., Krause, J., Briggs, A. W., Maricic, T., Stenzel, U., Kircher, M., Patterson, N., Li, H., Zhai, W., Fritz, M. H.-Y., Hansen, N. F., Durand, E. Y., Malaspinas, A.-S., Jensen, J. D., Marques-Bonet, T., Alkan, C., Prüfer, K., Meyer, M., Burbano, H. A., … Pääbo, S. (2010). A draft sequence of the neandertal genome. Science, 328 (5979), 710–722.

Grimwood, J., Gordon, L. A., Olsen, A., Terry, A., Schmutz, J., Lamerdin, J., Hellsten, U., Goodstein, D., Couronne, O., Tran-Gyamfi, M., Aerts, A., Altherr, M., Ashworth, L., Bajorek, E., Black, S., Branscomb, E., Caenepeel, S., Carrano, A., Caoile, C., … Lucas, S. M. (2004). The DNA sequence and biology of human chromosome 19. Nature, 428 (6982), 529–535.

Hamlin, J. A. P., Hibbins, M. S., & Moyle, L. C. (2020). Assessing biological factors affecting postspeciation introgression. Evol. Lett., 4 (2), 137–154.

Harris, K., & Nielsen, R. (2016). The genetic cost of neanderthal introgression. Genetics, 203 (2), 881–891.

Hedrick, P. W. (2013). Adaptive introgression in animals: Examples and comparison to new mutation and standing variation as sources of adaptive variation. Mol. Ecol., 22 (18), 4606– 4618.

Herrero, J., Muffato, M., Beal, K., Fitzgerald, S., Gordon, L., Pignatelli, M., Vilella, A. J., Searle, S. M. J., Amode, R., Brent, S., Spooner, W., Kulesha, E., Yates, A., & Flicek, P. (2016). Ensembl comparative genomics resources. Database, 2016.

Huerta-Sánchez, E., Jin, X., Asan Bianba, Z., Peter, B. M., Vinckenbosch, N., Liang, Y., Yi, X., He, M., Somel, M., Ni, P., Wang, B., Ou, X., Huasang Luosang, J., Cuo, Z. X. P., Li, K., Gao, G., Yin, Y., … Nielsen, R. (2014). Altitude adaptation in tibetans caused by introgression of denisovan-like DNA. Nature, 512 (7513), 194–197.

Jagoda, E., Xue, J. R., Reilly, S. K., Dannemann, M., Racimo, F., Huerta-Sanchez, E., Sankararaman, S., Kelso, J., Pagani, L., Sabeti, P. C., & Capellini, T. D. (2022). Detection of neanderthal adaptively introgressed genetic variants that modulate reporter gene expression in human immune cells. Mol. Biol. Evol., 39 (1).

Jones, M. R., Mills, L. S., Alves, P. C., Callahan, C. M., Alves, J. M., Lafferty, D. J. R., Jiggins, F. M., Jensen, J. D., Melo-Ferreira, J., & Good, J. M. (2018). Adaptive introgression underlies polymorphic seasonal camouflage in snowshoe hares. Science, 360 (6395), 1355– 1358.

Kim, B. Y., Huber, C. D., & Lohmueller, K. E. (2018). Deleterious variation shapes the genomic landscape of introgression. PLoS Genet., 14 (10), e1007741.

Li, H. (2011). A statistical framework for SNP calling, mutation discovery, association mapping and population genetical parameter estimation from sequencing data. Bioinformatics, 27 (21), 2987–2993.

Lopez Fang, L., Vecchyo, D. O.-D., McTavish, E. J., & Huerta-Sanchez, E. (2022). Leveraging shared ancestral variation to detect local introgression.

Mafessoni, F., Grote, S., de Filippo, C., Slon, V., Kolobova, K. A., Viola, B., Markin, S. V., Chintalapati, M., Peyrégne, S., Skov, L., Skoglund, P., Krivoshapkin, A. I., Derevianko, A. P., Meyer, M., Kelso, J., Peter, B., Prüfer, K., & Pääbo, S. (2020). A high-coverage neandertal genome from chagyrskaya cave. Proc. Natl. Acad. Sci. U. S. A., 117 (26), 15132– 15136.

Malinsky, M., Svardal, H., Tyers, A. M., Miska, E. A., Genner, M. J., Turner, G. F., & Durbin, R. (2018). Whole-genome sequences of malawi cichlids reveal multiple radiations interconnected by gene flow. Nat Ecol Evol, 2 (12), 1940–1955.

Martin, S. H., Davey, J. W., & Jiggins, C. D. (2014). Evaluating the use of ABBA–BABA statistics to locate introgressed loci. Mol. Biol. Evol., 32 (1), 244–257.

Meier, J. I., Marques, D. A., Mwaiko, S., Wagner, C. E., Excoffier, L., & Seehausen, O. (2017). Ancient hybridization fuels rapid cichlid fish adaptive radiations. Nat. Commun., 8, 14363.

Nelson, T. C., Stathos, A. M., Vanderpool, D. D., Finseth, F. R., Yuan, Y.-W., & Fishman, L. (2021). Ancient and recent introgression shape the evolutionary history of pollinator adaptation and speciation in a model monkeyflower radiation (mimulus section erythranthe). PLoS Genet., 17 (2), e1009095.

Patterson, N., Moorjani, P., Luo, Y., Mallick, S., Rohland, N., Zhan, Y., Genschoreck, T., Webster, T., & Reich, D. (2012). Ancient admixture in human history. Genetics, 192 (3), 1065–1093.

Pease, J. B., Haak, D. C., Hahn, M. W., & Moyle, L. C. (2016). Phylogenomics reveals three sources of adaptive variation during a rapid radiation. PLoS Biol., 14 (2), e1002379.

Pease, J. B., & Hahn, M. W. (2015). Detection and polarization of introgression in a Five-Taxon phylogeny. Syst. Biol., 64 (4), 651–662.

Pfeifer, B., & Kapan, D. D. (2019). Estimates of introgression as a function of pairwise distances. BMC Bioinformatics, 20 (1), 207.

Prüfer, K., de Filippo, C., Grote, S., Mafessoni, F., Korlević, P., Hajdinjak, M., Vernot, B., Skov, L., Hsieh, P., Peyrégne, S., Reher, D., Hopfe, C., Nagel, S., Maricic, T., Fu, Q., Theunert, C., Rogers, R., Skoglund, P., Chintalapati, M., … Pääbo, S. (2017). A high-coverage neandertal genome from vindija cave in croatia. Science, 358 (6363), 655–658.

Prüfer, K., Racimo, F., Patterson, N., Jay, F., Sankararaman, S., Sawyer, S., Heinze, A., Renaud, G., Sudmant, P. H., de Filippo, C., Li, H., Mallick, S., Dannemann, M., Fu, Q., Kircher, M., Kuhlwilm, M., Lachmann, M., Meyer, M., Ongyerth, M., … Pääbo, S. (2014). The complete genome sequence of a neanderthal from the altai mountains. Nature, 505 (7481), 43–49.

Racimo, F., Marnetto, D., & Huerta-Sánchez, E. (2016). Signatures of archaic adaptive introgression in Present-Day human populations. Mol. Biol. Evol., 34 (2), 296–317.

Ragsdale, A. P., & Gravel, S. (2019). Models of archaic admixture and recent history from two-locus statistics. PLoS Genet., 15 (6), e1008204.

Rheindt, F. E., & Edwards, S. V. (2011). Genetic introgression: An integral but neglected component of speciation in birds. Auk, 128 (4), 620–632.

Richards, E. J., & Martin, C. H. (2017). Adaptive introgression from distant caribbean islands contributed to the diversification of a microendemic adaptive radiation of trophic specialist pupfishes. PLoS Genet., 13 (8), e1006919.

Rogers, A. R., & Bohlender, R. J. (2015). Bias in estimators of archaic admixture. Theor. Popul. Biol., 100C, 63–78.

Stankowski, S., & Streisfeld, M. A. (2015). Introgressive hybridization facilitates adaptive divergence in a recent radiation of monkeyflowers. Proc. Biol. Sci., 282 (1814).

Suarez-Gonzalez, A., Lexer, C., & Cronk, Q. C. B. (2018). Adaptive introgression: A plant perspective. Biol. Lett., 14 (3).

Villanea, F. A., & Schraiber, J. G. (2019). Multiple episodes of interbreeding between neanderthal and modern humans. Nat Ecol Evol, 3 (1), 39–44.

Virtanen, P., Gommers, R., Oliphant, T. E., Haberland, M., Reddy, T., Cournapeau, D., Burovski, E., Peterson, P., Weckesser, W., Bright, J., van der Walt, S. J., Brett, M., Wilson, J., Millman, K. J., Mayorov, N., Nelson, A. R. J., Jones, E., Kern, R., Larson, E., … SciPy 1.0 Contributors. (2020). SciPy 1.0: Fundamental Algorithms for Scientific Computing in Python. Nature Methods, 17, 261–272. https://doi.org/10.1038/s41592-019-0686-2

Zhang, X., Witt, K. E., Bañuelos, M. M., Ko, A., Yuan, K., Xu, S., Nielsen, R., & Huerta-Sanchez, E. (2021). The history and evolution of the Denisovan-EPAS1 haplotype in tibetans. Proc. Natl. Acad. Sci. U. S. A., 118 (22).

