## Appendix for "The Utility of Ancestral and Derived Allele Sharing for Genome-Wide Inferences of Introgression"

### Appendix: Introgression Metrics Expectations and Equations

#### 1 Instantaneous Unidirectional Admixture Model of Introgression

Figure A1 graphically depicts the instantaneous unidirectional admixture (IUA) model of introgression, which assumes a three-taxon species tree—in Newick format: ((*P1*, *P2*), *P3*)—where the *P1* and *P2* populations diverged from an ancestral population *P12* at time $T_{P2}$ and where the *P3* and *P12* populations diverged from an ancestral population *P123* at time $T_{P3}$ (Green et al. 2010; Durand et al. 2011). Additionally, the IUA model assumes that a single introgression event took place at time $T_{GF}$, which can occur at any generation between $T_{P2}$ and generation zero, with a probability $f$ that a *P2* lineage traces its ancestry from the *P3* side of the phylogeny. Additionally, we assume that all populations have an equal and constant effective population size throughout the entire demographic history, an infinite sites mutation model, and that ancestral states are known. It should be noted that in practice one would also consider an additional taxon—in Newick format: (((*P1*, *P2*), *P3*), *O*)—where the *O* population is ancestral to the *P123* population, and is used as a putative outgroup to polarize ancestral states, since true ancestral states are rarely known. The IUA model was used to derive the expectations for the expected branch lengths that would generate an *ABBA*, *BABA*, *BAAA*, and *ABAA* site patterns, and we leverage these previously described expectations to derive the coalescent based expectations for the introgression metrics explored in this paper (Green et al. 2010; Durand et al. 2011; Lopez Fang et al. 2022).

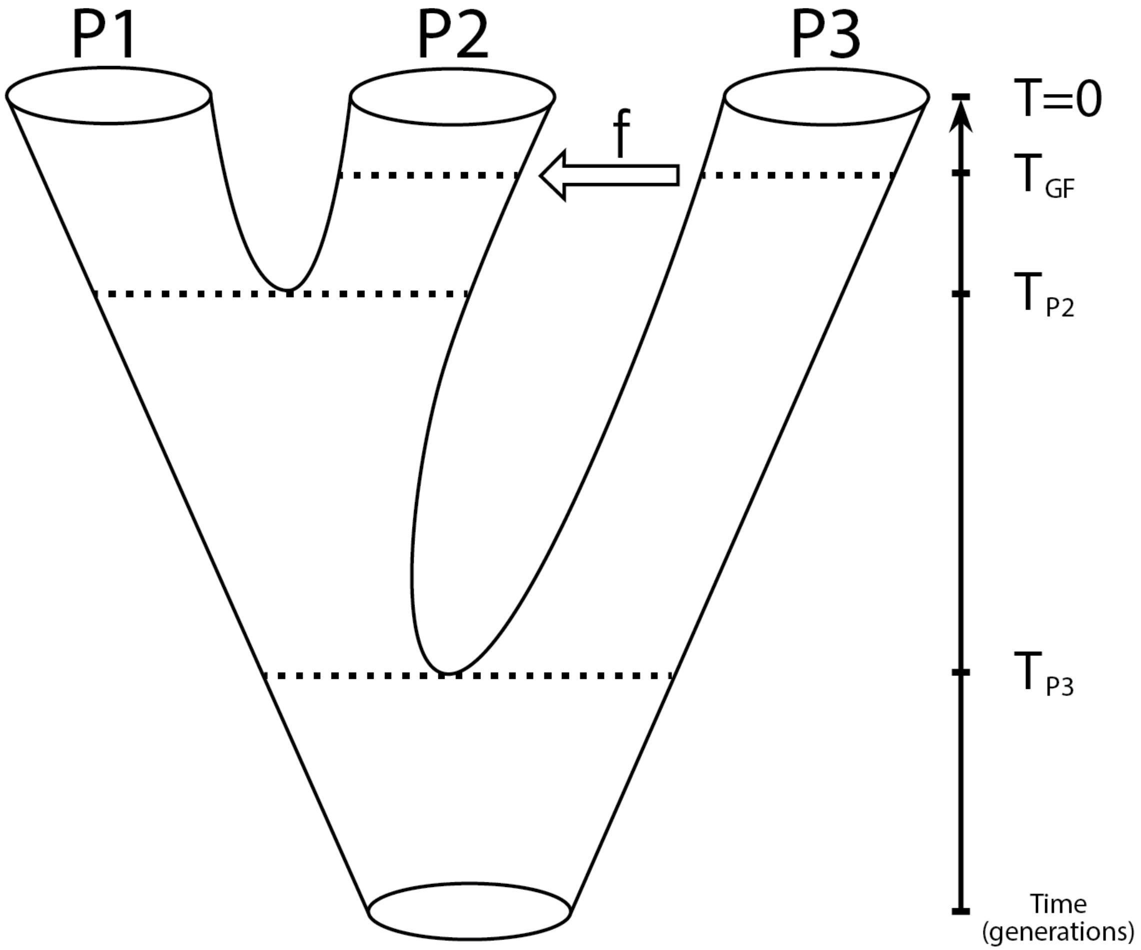


Figure A1: Instantaneous unidirectional admixture model (IUA) of introgression.

#### 2 Detection Metrics

The detection metrics explored in this paper tests for a significant excess of allele sharing between the donor and recipient populations. This is accomplished by using observed site patterns as a proxy for genealogical relationships between populations, however, it should be noted that the null hypothesis differs between detection metrics. Specifically, a statistically significant non-zero value of *Patterson's D* should be interpreted as evidence for rejecting an incomplete lineage sorting (ILS) only model, while statistically significant non-zero values of $D_{anc}$ and $D+$ should be interpreted as evidence for rejecting a model with no introgression (Green et al. 2010; Durand et al. 2011; Lopez Fang et al. 2022). This is due to the fact that in the absence of introgression it is only possible to generate *ABBA* and *BABA* site patterns via genealogical histories of ILS, while conversely the *BAAA* and *ABAA* site patterns may be generated via histories of ILS or LS (Green et al. 2010; Durand et al. 2011; Lopez Fang et al. 2022). For full derivations of *Patterson's D* reference *Durand et al. 2011* and for full derivations of $D_{anc}$ and $D+$ reference *Lopez-Fang et al. 2022.*

##### 2.1 Coalescent Expectations

$$D = \frac{(f) \cdot (T_{P3} - T_{GF})}{(1-f) \cdot \left[ {}^{4N}\!/_{3} \cdot e^{\frac{-(T_{P3}-T_{P2})}{2N}} \right] + (f) \cdot \left[ \left( {}^{4N}\!/_{3} \cdot e^{\frac{-(T_{P3}-T_{GF})}{2N}} \right) + (T_{P3} - T_{GF}) \right]} \tag{A1}$$

$$D_{anc} = \frac{(f) \cdot (T_{P3} - T_{GF})}{(1-f) \cdot \left[ 4N - \left( {}^{4N}\!/_{3} \cdot e^{\frac{-(T_{P3}-T_{P2})}{2N}} \right) + (2 \cdot T_{P2}) \right] + (f) \cdot \left[ (2N + T_{GF}) - \left( {}^{4N}\!/_{3} \cdot e^{\frac{-(T_{P3}-T_{GF})}{2N}} \right) + (2N + T_{P3}) \right]} \tag{A2}$$

$$D+ = \frac{(2) \cdot (f) \cdot (T_{P3} - T_{GF})}{(1-f) \cdot [4N + (2 \cdot T_{P2})] + (f) \cdot [4N + (2 \cdot T_{P3})]} \tag{A3}$$

##### 2.2 Derived Allele Frequency Equations

$$D = \frac{\sum_{i=1}^{n} (1-p_{i1}) p_{i2} p_{i3} (1-p_{iO}) - p_{i1}(1-p_{i2}) p_{i3} (1-p_{iO})}{\sum_{i=1}^{n} (1-p_{i1}) p_{i2} p_{i3} (1-p_{iO}) + p_{i1}(1-p_{i2}) p_{i3} (1-p_{iO})} \tag{A4}$$

$$D_{anc} = \frac{\sum_{i=1}^{n} p_{i1}(1-p_{i2})(1-p_{i3})(1-p_{iO}) - (1-p_{i1})p_{i2}(1-p_{i3})(1-p_{iO})}{\sum_{i=1}^{n} p_{i1}(1-p_{i2})(1-p_{i3})(1-p_{iO}) + (1-p_{i1})p_{i2}(1-p_{i3})(1-p_{iO})} \tag{A5}$$

$$D+ = \frac{\sum_{i=1}^{n}[(1-p_{i1})p_{i2}p_{i3}(1-p_{iO}) - p_{i1}(1-p_{i2})p_{i3}(1-p_{iO})] + [p_{i1}(1-p_{i2})(1-p_{i3})(1-p_{iO}) - (1-p_{i1})p_{i2}(1-p_{i3})(1-p_{iO})]}{\sum_{i=1}^{n}[(1-p_{i1})p_{i2}p_{i3}(1-p_{iO}) + p_{i1}(1-p_{i2})p_{i3}(1-p_{iO})] + [p_{i1}(1-p_{i2})(1-p_{i3})(1-p_{iO}) + (1-p_{i1})p_{i2}(1-p_{i3})(1-p_{iO})]} \tag{A6}$$

### 3 Quantification Metrics

The quantification metrics explored in this paper assesses the ratio of observed difference in allele sharing between *P3* and *P2* in the numerator and the expected differences assuming the entire genome was introgressed—i.e., complete homogenization of allele sharing—from *P3* to *P2* in the denominator. This is accomplished by replacing *P2* with *P3* in the assumed underlying species tree—i.e., in newick format: *(((P1, P3), P3,), O)*—when computing the denominator. Consequently, when we make the assumption that the entire genome was introgressed, in computing the denominator we assume that $f = 1$ and $T_{GF} = 0$.

#### 3.1 Coalescent Expectations

$$S\,(P1, P2, P3, O) = \mathbb{E}\,[\tau_{ABBA}] - \mathbb{E}\,[\tau_{BABA}] = (f) \cdot (T_{P3} - T_{GF}) \tag{A7a}$$

$$S\,(P1, P3, P3, O) = \mathbb{E}\,[\tau_{ABBA}] - \mathbb{E}\,[\tau_{BABA}] = T_{P3} + 2N \tag{A7b}$$

$$f_{hom} = \frac{S\,(P1, P2, P3, O)}{S\,(P1, P3, P3, O)} = \frac{(f) \cdot (T_{P3} - T_{GF})}{T_{P3} + 2N} \tag{A7c}$$

$$S_{ANC}\,(P1, P2, P3, O) = \mathbb{E}\,[\tau_{BAAA}] - \mathbb{E}\,[\tau_{ABAA}] = (f) \cdot (T_{P3} - T_{GF}) \tag{A8a}$$

$$S_{ANC}\,(P1, P3, P3, O) = \mathbb{E}\,[\tau_{BAAA}] - \mathbb{E}\,[\tau_{ABAA}] = T_{P3} + 2N \tag{A8b}$$

$$f_{anc} = \frac{S_{ANC}\,(P1, P2, P3, O)}{S_{ANC}\,(P1, P3, P3, O)} = \frac{(f) \cdot (T_{P3} - T_{GF})}{T_{P3} + 2N} \tag{A8c}$$

$$S+(P1,P2,P3,O)=(\mathbb{E}\left[\tau_{ABBA}\right]-\mathbb{E}\left[\tau_{BABA}\right])+(\mathbb{E}\left[\tau_{BAAA}\right]-\mathbb{E}\left[\tau_{ABAA}\right])=(2)\bullet(f)\bullet(T_{P3}-T_{GF}) \tag{A9a}$$

$$S+(P1,P3,P3,O)=(\mathbb{E}\left[\tau_{ABBA}\right]-\mathbb{E}\left[\tau_{BABA}\right])+(\mathbb{E}\left[\tau_{BAAA}\right]-\mathbb{E}\left[\tau_{ABAA}\right])=(2)\bullet(T_{P3}+2N) \tag{A9b}$$

$$f+=\frac{S+(P1,P2,P3,O)}{S+(P1,P3,P3,O)}=\frac{(f)\bullet(T_{P3}-T_{GF})}{T_{P3}+2N} \tag{A9c}$$

##### 3.2 Derived Allele Frequency Equations

$$f_{hom}=\frac{\sum_{i=1}^{n}(1-p_{i1})p_{i2}p_{i3}(1-p_{iO})-p_{i1}(1-p_{i2})p_{i3}(1-p_{iO})}{\sum_{i=1}^{n}(1-p_{i1})p_{i3}p_{i3}(1-p_{iO})-p_{i1}(1-p_{i3})p_{i3}(1-p_{iO})} \tag{A10}$$

$$f_{anc}=\frac{\sum_{i=1}^{n}p_{i1}(1-p_{i2})(1-p_{i3})(1-p_{iO})-(1-p_{i1})p_{i2}(1-p_{i3})(1-p_{iO})}{\sum_{i=1}^{n}p_{i1}(1-p_{i3})(1-p_{i3})(1-p_{iO})-(1-p_{i1})p_{i3}(1-p_{i3})(1-p_{iO})} \tag{A11}$$

$$f+=\frac{\sum_{i=1}^{n}[(1-p_{i1})p_{i2}p_{i3}(1-p_{iO})-p_{i1}(1-p_{i2})p_{i3}(1-p_{iO})]+[p_{i1}(1-p_{i2})(1-p_{i3})(1-p_{iO})-(1-p_{i1})p_{i2}(1-p_{i3})(1-p_{iO})]}{\sum_{i=1}^{n}[(1-p_{i1})p_{i3}p_{i3}(1-p_{iO})-p_{i1}(1-p_{i3})p_{i3}(1-p_{iO})]+[p_{i1}(1-p_{i3})(1-p_{i3})(1-p_{iO})-(1-p_{i1})p_{i3}(1-p_{i3})(1-p_{iO})]} \tag{A12}$$
