## Supplement for "The Utility of Ancestral and Derived Allele Sharing for Genome-Wide Inferences of Introgression"

### Supplementary Figures & Tables

### 1 Figures

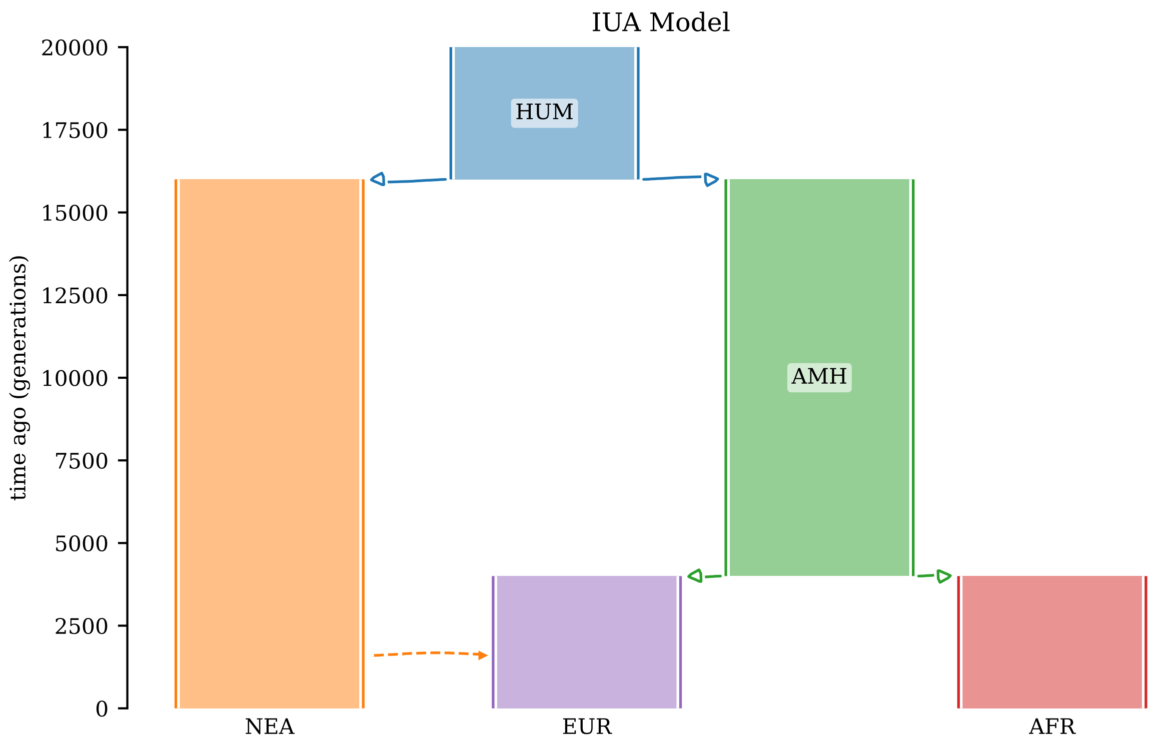


Figure S1: Instantaneous unidirectional admixture (IUA)—based on a simplified version of human evolution from *Racimo et. al., 2017*—used in this study. Solid lines represent divergence events and the dashed line represents a gene flow event.

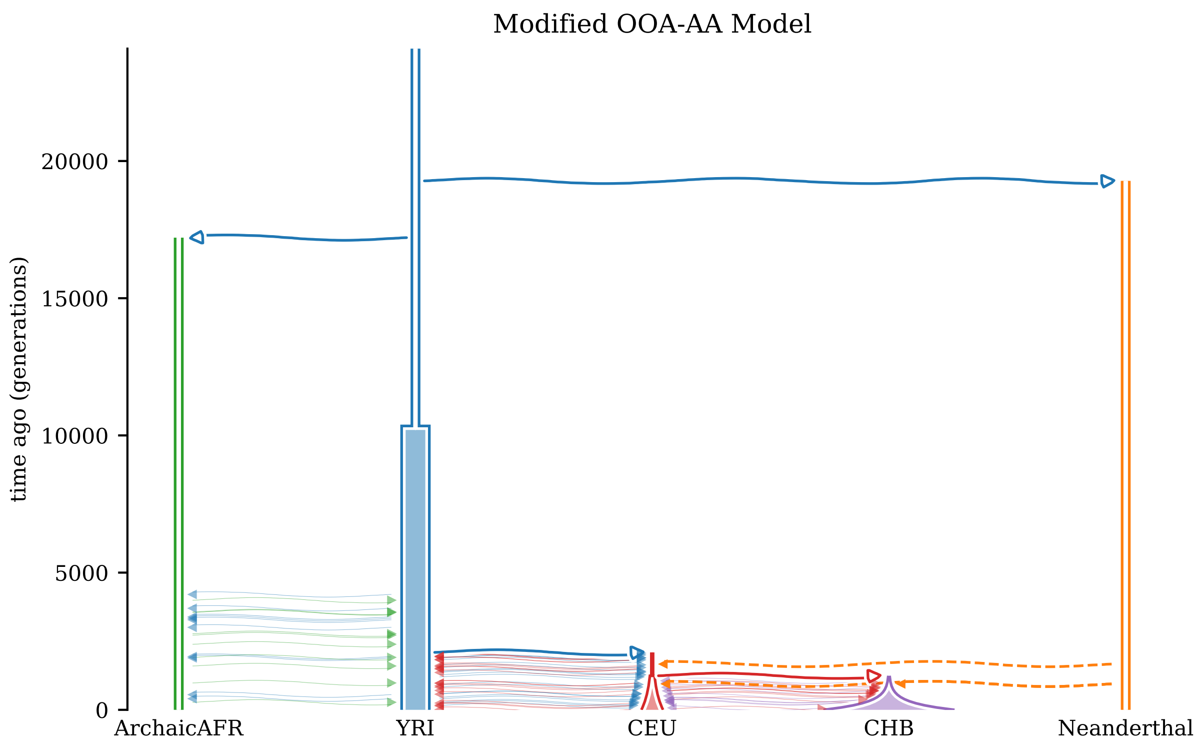


Figure S2: Realistic model of human demography—modified from *Ragsdale and Gravel 2019*—used in this study. Solid lines represent divergence events and dashed lines represents a gene flow events.

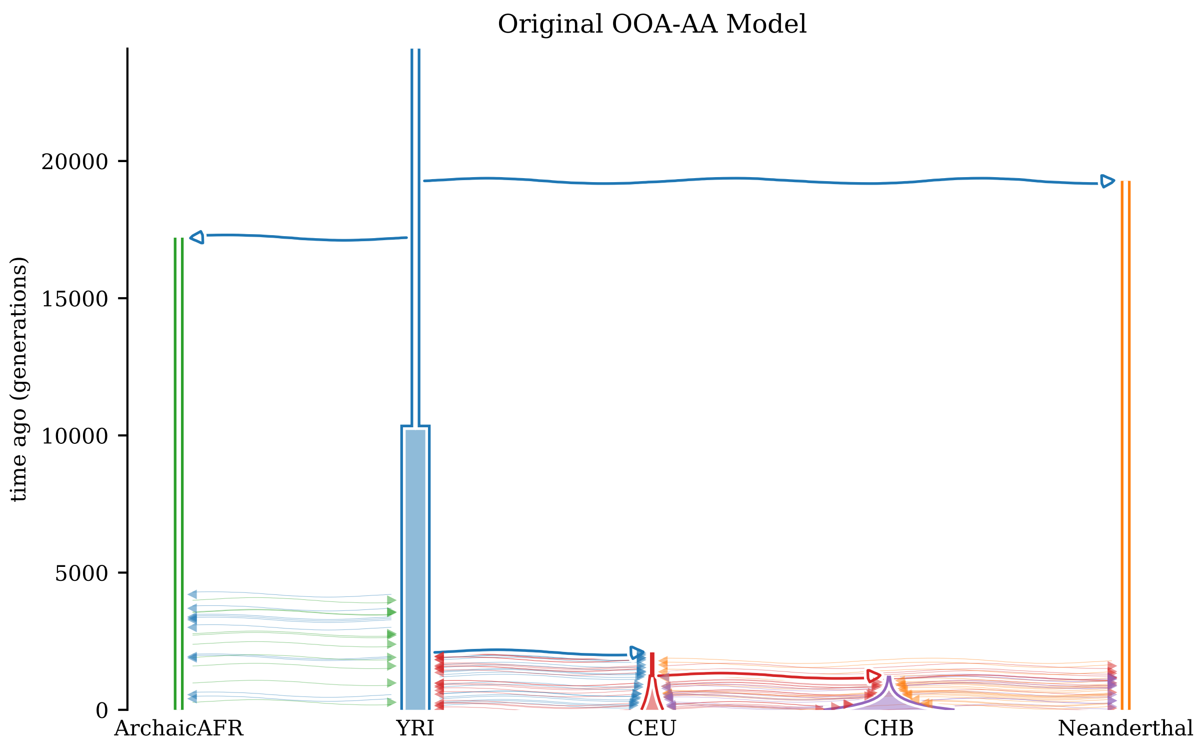


Figure S3: Realistic model of human demography—as originally specified in *Ragsdale and Gravel 2019*—used in this study. Solid lines represent divergence events and dashed lines represents a gene flow events.

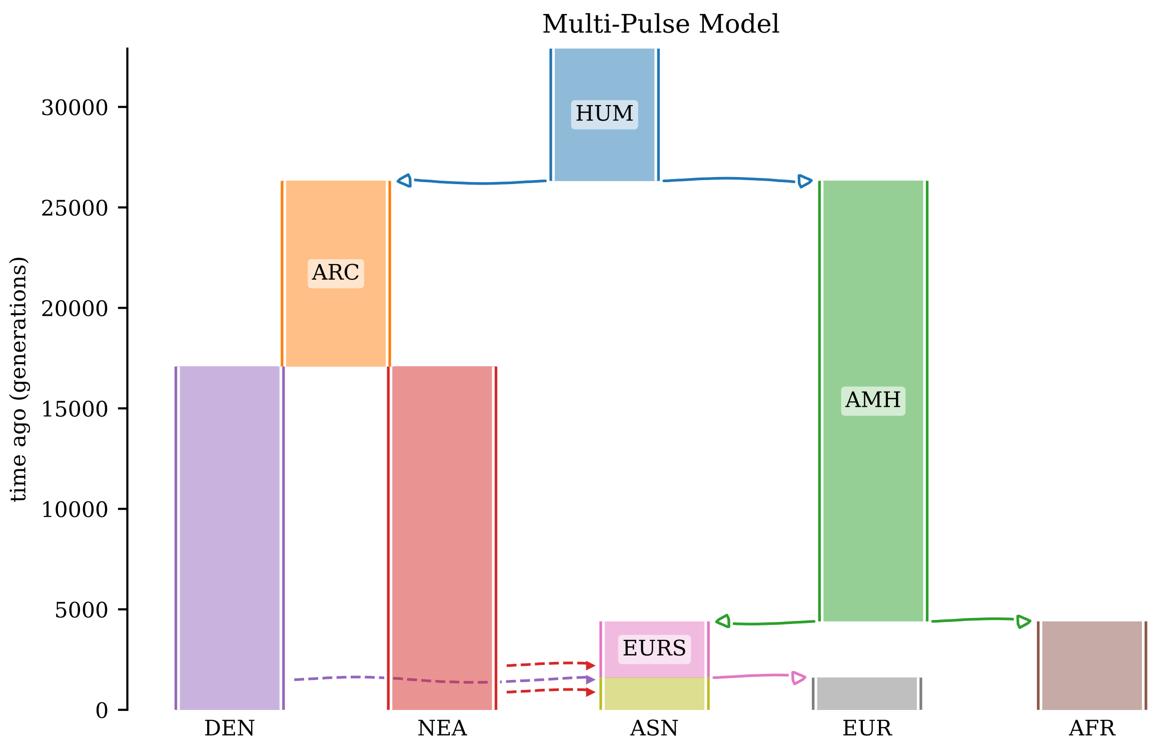


Figure S4: Multi-pulse model of archaic introgression used in this study. Solid lines represent divergence events and dashed lines represents a gene flow events.

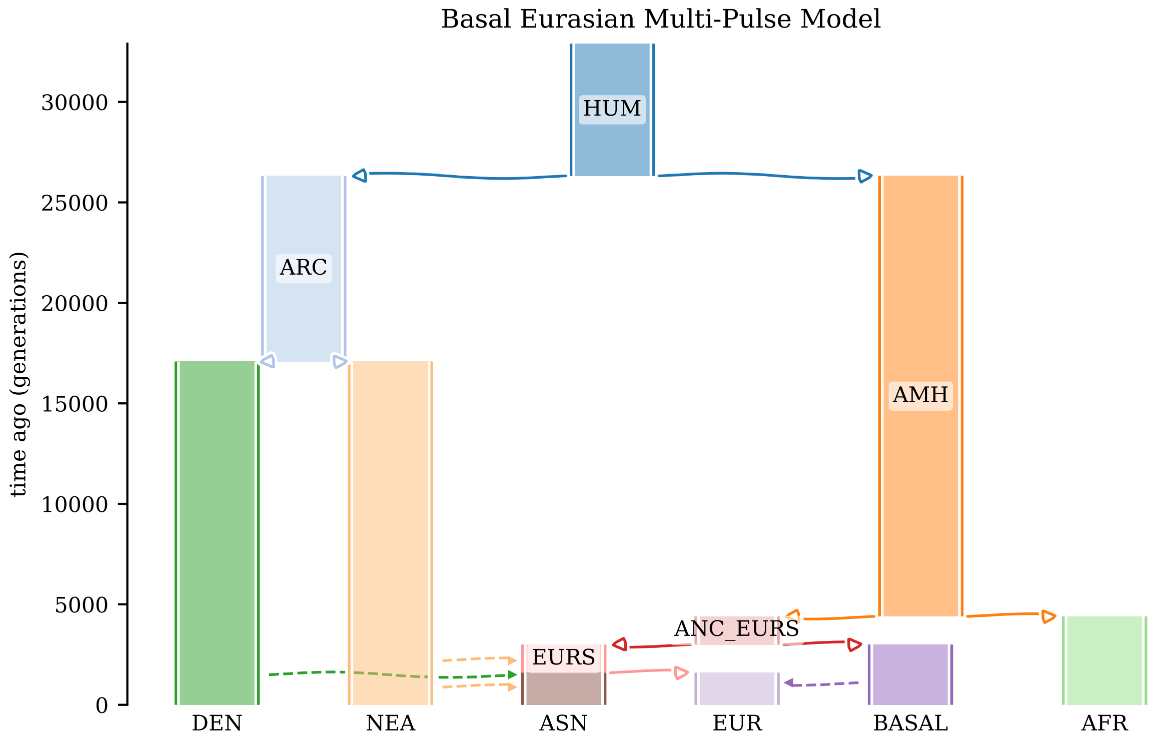


Figure S5: Multi-pulse model of archaic introgression with gene flow from a so called "Basal Eurasian" population used in this study. Solid lines represent divergence events and dashed lines represents a gene flow events.

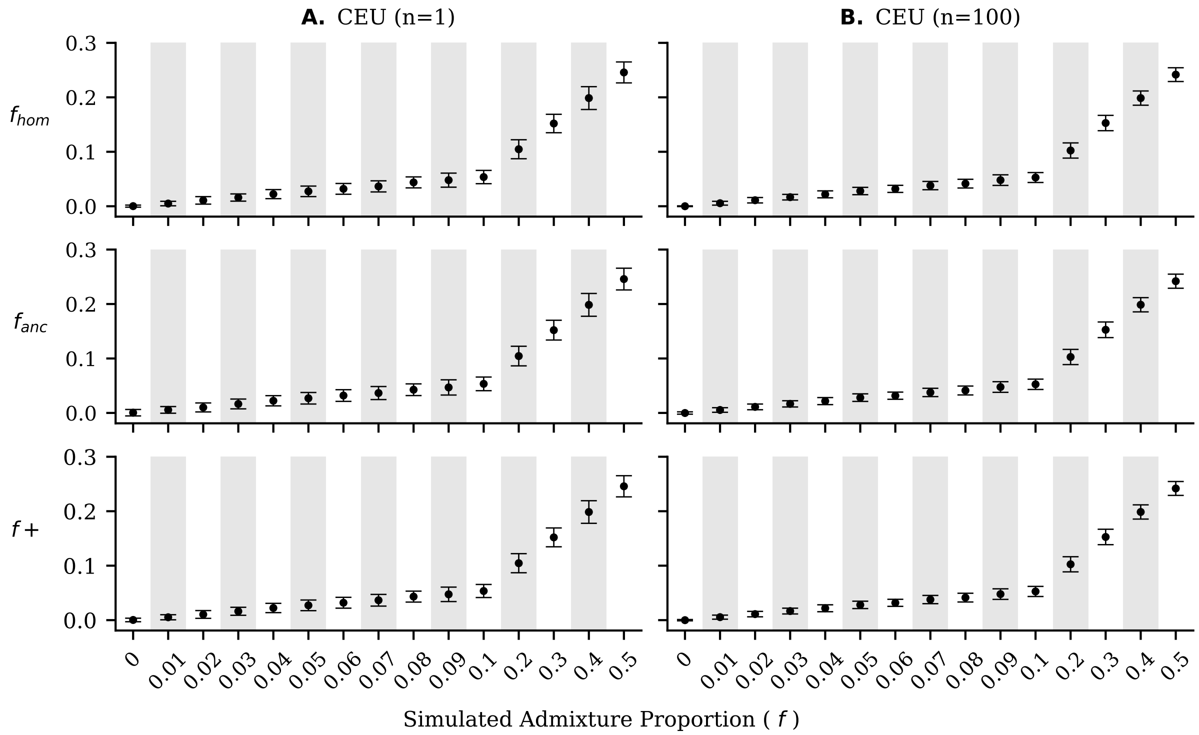


Figure S6: The mean (dots) and standard deviation (error bars) of $f_{hom}$, $f_{anc}$, and $f+$ for the CEU population in simulations under a more realistic model of human demography with a sampling scheme of n = 1 (panel **A**) and n = 100 (panel **B**) genomes from each of the potential recipient populations.

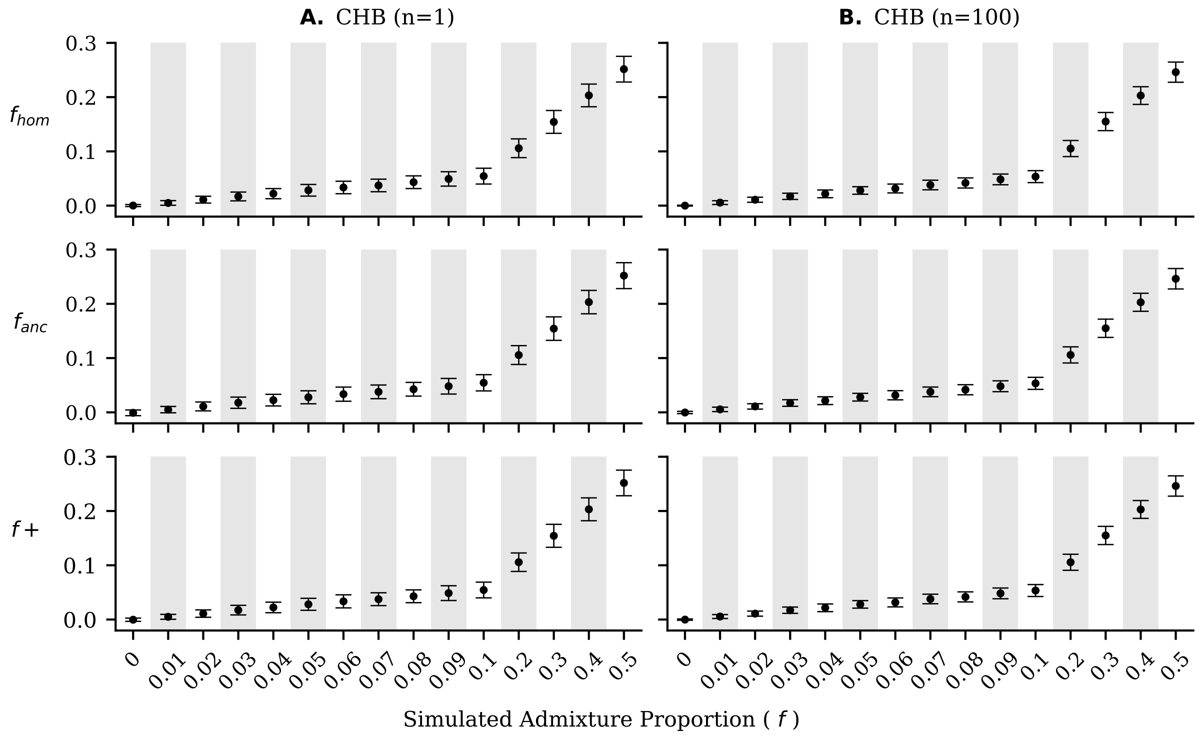


Figure S7: The mean (dots) and standard deviation (error bars) of $f_{hom}$, $f_{anc}$, and $f+$ for the CHB population in simulations under a more realistic model of human demography with a sampling scheme of n = 1 (panel **A**) and n = 100 (panel **B**) genomes from each of the potential recipient populations.

### 2 Tables

| | $n = 1$ | | | | | | | | $n = 100$ | | | | | | | |
|---|---|---|---|---|---|---|---|---|---|---|---|---|---|---|---|---|
| | $ABBA$ | | $BABA$ | | $BAAA$ | | $ABAA$ | | $ABBA$ | | $BABA$ | | $BAAA$ | | $ABAA$ | |
| | $\mu$ | $\sigma$ | $\mu$ | $\sigma$ | $\mu$ | $\sigma$ | $\mu$ | $\sigma$ | $\mu$ | $\sigma$ | $\mu$ | $\sigma$ | $\mu$ | $\sigma$ | $\mu$ | $\sigma$ |
| $f = 0$ | 5465.17 | 145.786 | 5495.06 | 174.939 | 30466.83 | 30466.83 | 30504.71 | 360.091 | 5503.459 | 99.246 | 5493.25 | 89.21 | 30521.818 | 30521.818 | 30510.263 | 173.3 |
| $f = 0.01$ | 5717.9 | 162.822 | 5482.68 | 150.028 | 30691.47 | 30691.47 | 30441.17 | 306.834 | 5683.277 | 99.461 | 5491.361 | 94.012 | 30703.991 | 30703.991 | 30501.073 | 174.717 |
| $f = 0.02$ | 5915.09 | 207.047 | 5486.46 | 134.769 | 30886.86 | 30886.86 | 30489.11 | 322.086 | 5918.106 | 116.926 | 5475.363 | 86.416 | 30892.775 | 30892.775 | 30469.369 | 165.639 |
| $f = 0.03$ | 6128.35 | 219.137 | 5464.9 | 151.926 | 31080.67 | 31080.67 | 30418.99 | 330.756 | 6109.743 | 133.018 | 5466.721 | 87.228 | 31066.999 | 31066.999 | 30430.308 | 152.027 |
| $f = 0.04$ | 6344.98 | 224.19 | 5457.72 | 153.123 | 31252.23 | 31252.23 | 30344.05 | 292.802 | 6322.632 | 157.614 | 5460.399 | 92.418 | 31270.053 | 31270.053 | 30412.161 | 148.245 |
| $f = 0.05$ | 6523.7 | 272.163 | 5467.93 | 165.331 | 31428.86 | 31428.86 | 30371.63 | 314.048 | 6525.706 | 162.207 | 5462.683 | 90.809 | 31430.519 | 31430.519 | 30377.307 | 169.344 |
| $f = 0.06$ | 6763.24 | 310.029 | 5447.03 | 151.423 | 31605.59 | 31605.59 | 30364.75 | 299.154 | 6731.675 | 168.588 | 5456.044 | 80.457 | 31613.554 | 31613.554 | 30330.141 | 184.834 |
| $f = 0.07$ | 6979.47 | 295.38 | 5444.42 | 152.392 | 31835.4 | 31835.4 | 30309.45 | 331.667 | 6948.888 | 189.516 | 5440.227 | 98.226 | 31807.404 | 31807.404 | 30296.78 | 177.464 |
| $f = 0.08$ | 7183.91 | 295.533 | 5442.11 | 154.242 | 32005.34 | 32005.34 | 30260.73 | 304.602 | 7162.842 | 183.305 | 5436.045 | 81.119 | 31959.277 | 31959.277 | 30256.61 | 166.038 |
| $f = 0.09$ | 7366.55 | 330.042 | 5408.98 | 146.531 | 32151.19 | 32151.19 | 30223.34 | 309.618 | 7397.239 | 207.577 | 5442.056 | 89.398 | 32212.754 | 32212.754 | 30258.873 | 175.291 |
| $f = 0.1$ | 7574.85 | 345.14 | 5438.46 | 162.725 | 32376.43 | 32376.43 | 30235.37 | 304.808 | 7602.723 | 216.035 | 5420.829 | 80.242 | 32383.078 | 32383.078 | 30219.948 | 164.952 |
| $f = 0.2$ | 9680.98 | 392.051 | 5365.18 | 154.698 | 34201.91 | 34201.91 | 29883.61 | 295.968 | 9737.691 | 278.78 | 5373.982 | 80.579 | 34282.719 | 34282.719 | 29940.206 | 160.401 |
| $f = 0.3$ | 11775.45 | 458.298 | 5312.79 | 162.303 | 36121.95 | 36121.95 | 29622.88 | 286.636 | 11820.454 | 323.273 | 5303.895 | 71.677 | 36156.464 | 36156.464 | 29614.956 | 156.411 |
| $f = 0.4$ | 13808.99 | 545.123 | 5239.44 | 146.509 | 37905.85 | 37905.85 | 29348.76 | 321.16 | 13890.801 | 306.937 | 5231.154 | 82.028 | 37961.395 | 37961.395 | 29310.817 | 139.544 |
| $f = 0.5$ | 15888.34 | 571.99 | 5169.04 | 160.197 | 39733.58 | 39733.58 | 28999.17 | 338.799 | 15973.043 | 326.765 | 5178.227 | 71.947 | 39825.797 | 39825.797 | 29028.167 | 163.446 |

Table S1: The mean ($\mu$) and standard deviation ($\sigma$) of each allele sharing site pattern between donor and recipient populations from 100 replicate simulations under an IUA model of introgression (see Figure S1) per admixture portion ($f$).

| | $n = 1$ | | | | | | | | $n = 100$ | | | | | | | |
|---|---|---|---|---|---|---|---|---|---|---|---|---|---|---|---|---|
| | $ABBA$ | | $BABA$ | | $BAAA$ | | $ABAA$ | | $ABBA$ | | $BABA$ | | $BAAA$ | | $ABAA$ | |
| | $\mu$ | $\sigma$ | $\mu$ | $\sigma$ | $\mu$ | $\sigma$ | $\mu$ | $\sigma$ | $\mu$ | $\sigma$ | $\mu$ | $\sigma$ | $\mu$ | $\sigma$ | $\mu$ | $\sigma$ |
| $f = 0$ | 930.51 | 64.016 | 922.82 | 76.896 | 21145.85 | 21145.85 | 21128.37 | 422.106 | 925.594 | 38.574 | 924.856 | 36.317 | 21172.813 | 21172.813 | 21178.431 | 177.082 |
| $f = 0.01$ | 1146.04 | 164.083 | 950.32 | 73.325 | 21308.29 | 21308.29 | 21085.87 | 355.821 | 1161.299 | 151.017 | 941.615 | 37.937 | 21347.06 | 21347.06 | 21132.82 | 173.405 |
| $f = 0.02$ | 1388.87 | 270.689 | 962.04 | 82.514 | 21449.92 | 21449.92 | 21049.05 | 355.924 | 1395.229 | 216.809 | 958.382 | 45.624 | 21492.765 | 21492.765 | 21052.189 | 155.638 |
| $f = 0.03$ | 1605.62 | 278.711 | 971.92 | 91.535 | 21665.74 | 21665.74 | 21010.99 | 375.695 | 1641.213 | 222.986 | 981.813 | 45.752 | 21649.188 | 21649.188 | 20988.938 | 165.208 |
| $f = 0.04$ | 1875.73 | 353.863 | 995.04 | 105.142 | 21745.71 | 21745.71 | 20860.5 | 402.936 | 1858.153 | 278.112 | 993.388 | 52.834 | 21770.682 | 21770.682 | 20909.659 | 175.301 |
| $f = 0.05$ | 2098.58 | 416.521 | 1016.95 | 105.916 | 21873.99 | 21873.99 | 20808.6 | 339.641 | 2122.177 | 302.454 | 1018.54 | 59.959 | 21935.686 | 21935.686 | 20825.567 | 154.587 |
| $f = 0.06$ | 2301.22 | 401.378 | 1038.42 | 103.689 | 22059.52 | 22059.52 | 20795.22 | 421.01 | 2288.618 | 281.98 | 1028.601 | 58.389 | 22033.893 | 22033.893 | 20783.524 | 185.324 |
| $f = 0.07$ | 2487.48 | 422.636 | 1047.17 | 118.077 | 22214.41 | 22214.41 | 20770.76 | 392.179 | 2553.619 | 343.669 | 1055.63 | 69.962 | 22227.236 | 22227.236 | 20740.811 | 165.605 |
| $f = 0.08$ | 2799.79 | 420.128 | 1070.04 | 124.241 | 22376.21 | 22376.21 | 20692.76 | 387.733 | 2691.676 | 343.483 | 1054.78 | 59.835 | 22316.171 | 22316.171 | 20689.874 | 163.512 |
| $f = 0.09$ | 2980.61 | 522.222 | 1092.56 | 104.837 | 22497.67 | 22497.67 | 20650.51 | 408.156 | 2985.191 | 417.713 | 1092.76 | 67.888 | 22529.64 | 22529.64 | 20651.583 | 186.357 |
| $f = 0.1$ | 3211.27 | 489.95 | 1094.5 | 133.077 | 22680.11 | 22680.11 | 20573.87 | 368.508 | 3166.78 | 384.45 | 1088.685 | 66.258 | 22644.868 | 22644.868 | 20569.972 | 207.883 |
| $f = 0.2$ | 5323.67 | 666.195 | 1224.73 | 143.135 | 24080.39 | 24080.39 | 19992.0 | 371.823 | 5244.473 | 586.663 | 1235.684 | 84.679 | 24003.713 | 24003.713 | 19979.526 | 240.193 |
| $f = 0.3$ | 7260.47 | 651.261 | 1355.05 | 154.912 | 25267.78 | 25267.78 | 19359.9 | 421.043 | 7290.452 | 568.385 | 1354.797 | 84.671 | 25313.556 | 25313.556 | 19375.832 | 250.17 |
| $f = 0.4$ | 9100.43 | 769.168 | 1435.45 | 184.026 | 26417.33 | 26417.33 | 18760.38 | 434.491 | 9092.109 | 514.219 | 1429.62 | 81.405 | 26440.301 | 26440.301 | 18771.175 | 207.241 |
| $f = 0.5$ | 10884.17 | 697.749 | 1440.74 | 166.488 | 27593.89 | 27593.89 | 18151.78 | 410.824 | 10739.418 | 482.943 | 1478.097 | 80.529 | 27465.756 | 27465.756 | 18190.12 | 216.318 |
| Original Model | 1481.11 | 276.914 | 1018.94 | 80.92 | 21270.2 | 21270.2 | 20808.1 | 353.2 | 1533.123 | 202.111 | 1025.721 | 52.224 | 21370.501 | 21370.501 | 20870.248 | 198.149 |

Table S2: The mean ($\mu$) and standard deviation ($\sigma$) of each allele sharing site pattern between the Neanderthal and CEU populations from 100 replicate simulations under a realistic model of human demography (see Figure S2) per admixture portion ($f$). Note that "Original Model" corresponds to the results using the original model from *Ragsdale and Gravel 2019* (see Figure S3).

| | $n = 1$ | | | | | | | | $n = 100$ | | | | | | | |
|---|---|---|---|---|---|---|---|---|---|---|---|---|---|---|---|---|
| | $ABBA$ | | $BABA$ | | $BAAA$ | | $ABAA$ | | $ABBA$ | | $BABA$ | | $BAAA$ | | $ABAA$ | |
| | $\mu$ | $\sigma$ | $\mu$ | $\sigma$ | $\mu$ | $\sigma$ | $\mu$ | $\sigma$ | $\mu$ | $\sigma$ | $\mu$ | $\sigma$ | $\mu$ | $\sigma$ | $\mu$ | $\sigma$ |
| $f = 0$ | 934.2 | 69.582 | 927.17 | 70.361 | 21182.93 | 21182.93 | 21217.99 | 431.995 | 924.622 | 39.63 | 925.532 | 37.311 | 21185.397 | 21185.397 | 21198.106 | 180.018 |
| $f = 0.01$ | 1150.93 | 168.358 | 953.36 | 71.185 | 21321.98 | 21321.98 | 21124.25 | 398.96 | 1162.986 | 146.833 | 943.296 | 39.222 | 21369.843 | 21369.843 | 21153.123 | 167.137 |
| $f = 0.02$ | 1400.73 | 250.677 | 963.89 | 82.524 | 21485.21 | 21485.21 | 21053.76 | 375.359 | 1391.294 | 203.48 | 961.332 | 46.067 | 21515.224 | 21515.224 | 21083.225 | 164.239 |
| $f = 0.03$ | 1640.96 | 325.808 | 971.01 | 92.789 | 21691.52 | 21691.52 | 20988.08 | 393.487 | 1661.809 | 246.33 | 982.643 | 44.592 | 21675.661 | 21675.661 | 20995.167 | 174.373 |
| $f = 0.04$ | 1877.06 | 370.903 | 1003.47 | 103.632 | 21773.5 | 21773.5 | 20884.64 | 425.134 | 1854.788 | 298.396 | 994.304 | 52.943 | 21778.46 | 21778.46 | 20928.016 | 188.018 |
| $f = 0.05$ | 2139.36 | 445.71 | 1020.34 | 106.041 | 21936.67 | 21936.67 | 20840.67 | 390.708 | 2126.662 | 301.511 | 1022.295 | 60.3 | 21957.289 | 21957.289 | 20854.561 | 183.226 |
| $f = 0.06$ | 2354.81 | 449.712 | 1031.17 | 110.067 | 22159.67 | 22159.67 | 20833.92 | 360.285 | 2277.733 | 348.574 | 1029.234 | 59.629 | 22041.557 | 22041.557 | 20798.666 | 197.698 |
| $f = 0.07$ | 2517.41 | 469.168 | 1045.95 | 115.556 | 22258.9 | 22258.9 | 20766.76 | 409.831 | 2559.627 | 381.779 | 1056.793 | 71.114 | 22247.489 | 22247.489 | 20756.667 | 199.83 |
| $f = 0.08$ | 2783.1 | 459.444 | 1076.98 | 133.706 | 22371.31 | 22371.31 | 20690.94 | 404.079 | 2704.243 | 384.563 | 1056.106 | 61.491 | 22342.556 | 22342.556 | 20699.08 | 181.359 |
| $f = 0.09$ | 3028.03 | 534.741 | 1087.47 | 113.787 | 22528.24 | 22528.24 | 20632.18 | 412.798 | 3002.136 | 416.357 | 1094.909 | 69.325 | 22560.492 | 22560.492 | 20664.256 | 201.644 |
| $f = 0.1$ | 3240.08 | 577.808 | 1092.25 | 128.416 | 22731.33 | 22731.33 | 20581.91 | 417.812 | 3195.566 | 448.746 | 1087.617 | 66.507 | 22682.428 | 22682.428 | 20575.972 | 221.881 |
| $f = 0.2$ | 5369.94 | 656.301 | 1227.8 | 145.219 | 24104.51 | 24104.51 | 19971.27 | 346.689 | 5352.274 | 610.597 | 1230.984 | 83.52 | 24090.155 | 24090.155 | 19945.377 | 258.798 |
| $f = 0.3$ | 7340.65 | 800.498 | 1342.61 | 144.623 | 25344.41 | 25344.41 | 19348.08 | 426.819 | 7375.947 | 657.394 | 1355.759 | 90.367 | 25385.743 | 25385.743 | 19360.986 | 285.147 |
| $f = 0.4$ | 9262.11 | 774.945 | 1421.74 | 169.275 | 26558.62 | 26558.62 | 18719.97 | 448.514 | 9253.655 | 626.236 | 1425.363 | 84.504 | 26559.397 | 26559.397 | 18728.226 | 251.869 |
| $f = 0.5$ | 11096.05 | 874.243 | 1429.86 | 178.433 | 27748.53 | 27748.53 | 18069.86 | 431.316 | 10898.188 | 696.852 | 1470.569 | 82.449 | 27582.945 | 27582.945 | 18143.087 | 298.144 |
| Original Model | 1512.69 | 316.969 | 1015.74 | 81.989 | 21342.76 | 21342.76 | 20820.99 | 336.052 | 1540.101 | 193.612 | 1021.449 | 48.881 | 21385.526 | 21385.526 | 20865.97 | 206.672 |

Table S3: The mean ($\mu$) and standard deviation ($\sigma$) of each allele sharing site pattern between the Neanderthal and CHB populations from 100 replicate simulations under a realistic model of human demography (see Figure S2) per admixture portion ($f$). Note that "Original Model" corresponds to the results using the original model from *Ragsdale and Gravel 2019* (see Figure S3).

| | $n = 1$ | | | $n = 100$ | | |
|---|---|---|---|---|---|---|
| | $ABBA - BABA$ | $BAAA - ABAA$ | $p - value$ | $ABBA - BABA$ | $BAAA - ABAA$ | $p - value$ |
| $f = 0$ | -29.89 | -37.88 | 0.487 | 10.209 | 11.556 | 0.495 |
| $f = 0.01$ | 235.22 | 250.3 | 0.475 | 191.916 | 202.918 | 0.459 |
| $f = 0.02$ | 428.63 | 397.75 | 0.459 | 442.743 | 423.406 | 0.438 |
| $f = 0.03$ | 663.45 | 661.68 | 0.497 | 643.022 | 636.691 | 0.477 |
| $f = 0.04$ | 887.26 | 908.18 | 0.467 | 862.233 | 857.893 | 0.485 |
| $f = 0.05$ | 1055.77 | 1057.23 | 0.498 | 1063.024 | 1053.212 | 0.464 |
| $f = 0.06$ | 1316.21 | 1240.84 | 0.395 | 1275.631 | 1283.414 | 0.471 |
| $f = 0.07$ | 1535.05 | 1525.95 | 0.486 | 1508.661 | 1510.623 | 0.493 |
| $f = 0.08$ | 1741.8 | 1744.61 | 0.496 | 1726.797 | 1702.667 | 0.413 |
| $f = 0.09$ | 1957.57 | 1927.85 | 0.456 | 1955.183 | 1953.881 | 0.495 |
| $f = 0.1$ | 2136.39 | 2141.06 | 0.494 | 2181.894 | 2163.13 | 0.434 |
| $f = 0.2$ | 4315.8 | 4318.3 | 0.496 | 4363.709 | 4342.513 | 0.434 |
| $f = 0.3$ | 6462.66 | 6499.07 | 0.448 | 6516.559 | 6541.508 | 0.426 |
| $f = 0.4$ | 8569.55 | 8557.09 | 0.484 | 8659.648 | 8650.578 | 0.472 |
| $f = 0.5$ | 10719.3 | 10734.41 | 0.479 | 10794.816 | 10797.63 | 0.493 |

Table S4: Assessing if $(ABBA - BABA) - (BAAA - ABAA)$ significantly differs from zero in simulations under an IUA model of introgression (see Figure S1). The rows correspond to the simulated admixture proportions, the columns represent the mean site pattern difference across 100 replicate simulations and the corresponding *p-value* to assess if $(ABBA - BABA) - (BAAA - ABAA)$ significantly differs from zero. After accounting for multiple-comparisons using the Bonferroni correction, a *p-value* less than 0.00333 is considered significant.

| | $n = 1$ | | | $n = 100$ | | |
|---|---|---|---|---|---|---|
| | $ABBA - BABA$ | $BAAA - ABAA$ | $p - value$ | $ABBA - BABA$ | $BAAA - ABAA$ | $p - value$ |
| $f = 0$ | 7.69 | 17.48 | 0.483 | 0.738 | -5.618 | 0.469 |
| $f = 0.01$ | 195.72 | 222.42 | 0.443 | 219.684 | 214.24 | 0.472 |
| $f = 0.02$ | 426.83 | 400.87 | 0.45 | 436.847 | 440.576 | 0.483 |
| $f = 0.03$ | 633.7 | 654.75 | 0.464 | 659.4 | 660.25 | 0.496 |
| $f = 0.04$ | 880.69 | 885.21 | 0.491 | 864.765 | 861.023 | 0.48 |
| $f = 0.05$ | 1081.63 | 1065.39 | 0.469 | 1103.637 | 1110.12 | 0.466 |
| $f = 0.06$ | 1262.8 | 1264.3 | 0.497 | 1260.017 | 1250.369 | 0.45 |
| $f = 0.07$ | 1440.31 | 1443.65 | 0.494 | 1497.99 | 1486.425 | 0.435 |
| $f = 0.08$ | 1729.75 | 1683.45 | 0.416 | 1636.895 | 1626.297 | 0.442 |
| $f = 0.09$ | 1888.05 | 1847.16 | 0.413 | 1892.432 | 1878.057 | 0.426 |
| $f = 0.1$ | 2116.77 | 2106.24 | 0.482 | 2078.096 | 2074.896 | 0.483 |
| $f = 0.2$ | 4098.94 | 4088.39 | 0.48 | 4008.79 | 4024.187 | 0.428 |
| $f = 0.3$ | 5905.42 | 5907.88 | 0.496 | 5935.655 | 5937.724 | 0.49 |
| $f = 0.4$ | 7664.98 | 7656.95 | 0.486 | 7662.49 | 7669.126 | 0.47 |
| $f = 0.5$ | 9443.43 | 9442.11 | 0.498 | 9261.32 | 9275.636 | 0.441 |
| Original Model | 462.17 | 462.1 | 0.5 | 507.402 | 500.254 | 0.469 |

Table S5: Assessing if *(ABBA − BABA) − (BAAA − ABAA)* significantly differs from zero in the CEU population under realistic simulations of human demography (see Figure S2). The rows correspond to the simulated admixture proportions, the columns represent the mean site pattern difference across 100 replicate simulations and the corresponding *p-value* to assess if *(ABBA − BABA) − (BAAA − ABAA)* significantly differs from zero. After accounting for multiple-comparisons using the Bonferroni correction, a *p-value* less than 0.003125 is considered significant. Note that "Original Model" corresponds to the results using the original model from *Ragsdale and Gravel 2019* (see Figure S3).

| | $n=1$ | | | $n=100$ | | |
|---|---|---|---|---|---|---|
| | $ABBA-BABA$ | $BAAA-ABAA$ | $p-value$ | $ABBA-BABA$ | $BAAA-ABAA$ | $p-value$ |
| $f=0$ | 7.03 | -35.06 | 0.418 | -0.909 | -12.709 | 0.442 |
| $f=0.01$ | 197.57 | 197.73 | 0.5 | 219.69 | 216.72 | 0.485 |
| $f=0.02$ | 436.84 | 431.45 | 0.49 | 429.962 | 432.0 | 0.491 |
| $f=0.03$ | 669.95 | 703.44 | 0.443 | 679.165 | 680.495 | 0.494 |
| $f=0.04$ | 873.59 | 888.86 | 0.47 | 860.484 | 850.444 | 0.45 |
| $f=0.05$ | 1119.02 | 1096.0 | 0.457 | 1104.367 | 1102.728 | 0.492 |
| $f=0.06$ | 1323.64 | 1325.75 | 0.496 | 1248.5 | 1242.891 | 0.472 |
| $f=0.07$ | 1471.46 | 1492.14 | 0.461 | 1502.834 | 1490.822 | 0.444 |
| $f=0.08$ | 1706.12 | 1680.37 | 0.459 | 1648.137 | 1643.475 | 0.478 |
| $f=0.09$ | 1940.56 | 1896.06 | 0.41 | 1907.226 | 1896.236 | 0.444 |
| $f=0.1$ | 2147.83 | 2149.42 | 0.497 | 2107.949 | 2106.456 | 0.492 |
| $f=0.2$ | 4142.14 | 4133.24 | 0.486 | 4121.289 | 4144.779 | 0.403 |
| $f=0.3$ | 5998.04 | 5996.33 | 0.497 | 6020.188 | 6024.757 | 0.48 |
| $f=0.4$ | 7840.37 | 7838.65 | 0.497 | 7828.291 | 7831.171 | 0.489 |
| $f=0.5$ | 9666.19 | 9678.67 | 0.481 | 9427.619 | 9439.857 | 0.45 |
| Original Model | 496.95 | 521.77 | 0.451 | 518.652 | 519.556 | 0.496 |

Table S6: Assessing if *(ABBA − BABA)* − *(BAAA − ABAA)* significantly differs from zero in the CHB population under realistic simulations of human demography (see Figure S2). The rows correspond to the simulated admixture proportions, the columns represent the mean site pattern difference across 100 replicate simulations and the corresponding *p-value* to assess if *(ABBA − BABA)* − *(BAAA − ABAA)* significantly differs from zero. After accounting for multiple-comparisons using the Bonferroni correction, a *p-value* less than 0.003125 is considered significant. Note that "Original Model" corresponds to the results using the original model from *Ragsdale and Gravel 2019* (see Figure S3).

| | $n=1$ | | | $n=100$ | | |
|---|---|---|---|---|---|---|
| | $D$ | $D_{anc}$ | $D+$ | $D$ | $D_{anc}$ | $D+$ |
| $f=0$ | 0.13 | 0.07 | 0.08 | 0.11 | 0.08 | 0.1 |
| $f=0.01$ | 0.34 | 0.27 | 0.34 | 0.66 | 0.5 | 0.6 |
| $f=0.02$ | 0.56 | 0.39 | 0.52 | 0.98 | 0.88 | 0.96 |
| $f=0.03$ | 0.87 | 0.63 | 0.86 | 1.0 | 0.98 | 1.0 |
| $f=0.04$ | 0.96 | 0.89 | 0.95 | 1.0 | 1.0 | 1.0 |
| $f=0.05$ | 0.99 | 0.92 | 0.99 | 1.0 | 1.0 | 1.0 |
| $f=0.06$ | 1.0 | 0.95 | 1.0 | 1.0 | 1.0 | 1.0 |
| $f=0.07$ | 1.0 | 0.97 | 1.0 | 1.0 | 1.0 | 1.0 |
| $f=0.08$ | 1.0 | 1.0 | 1.0 | 1.0 | 1.0 | 1.0 |
| $f=0.09$ | 1.0 | 1.0 | 1.0 | 1.0 | 1.0 | 1.0 |
| $f=0.1$ | 1.0 | 1.0 | 1.0 | 1.0 | 1.0 | 1.0 |
| $f=0.2$ | 1.0 | 1.0 | 1.0 | 1.0 | 1.0 | 1.0 |
| $f=0.3$ | 1.0 | 1.0 | 1.0 | 1.0 | 1.0 | 1.0 |
| $f=0.4$ | 1.0 | 1.0 | 1.0 | 1.0 | 1.0 | 1.0 |
| $f=0.5$ | 1.0 | 1.0 | 1.0 | 1.0 | 1.0 | 1.0 |

Table S7: Statistical power of $D$, $D_{anc}$, and $D+$ to detect introgression in simulations under an IUA model of introgression (see Figure S1). The rows correspond to the simulated admixture proportions and the columns represent the proportion of statistically significant replicates from zero at $p$-$value<0.05$ out of 100 replicate simulations.

| | $n = 1$ | | | $n = 100$ | | |
|---|---|---|---|---|---|---|
| | $D$ | $D_{anc}$ | $D+$ | $D$ | $D_{anc}$ | $D+$ |
| $f = 0$ | 0.13 | 0.11 | 0.13 | 0.0 | 0.0 | 0.0 |
| $f = 0.01$ | 0.54 | 0.25 | 0.44 | 0.6 | 0.2 | 0.39 |
| $f = 0.02$ | 0.87 | 0.46 | 0.66 | 1.0 | 0.54 | 0.85 |
| $f = 0.03$ | 1.0 | 0.78 | 0.89 | 1.0 | 0.9 | 1.0 |
| $f = 0.04$ | 1.0 | 0.88 | 0.94 | 1.0 | 0.98 | 1.0 |
| $f = 0.05$ | 1.0 | 0.93 | 1.0 | 1.0 | 1.0 | 1.0 |
| $f = 0.06$ | 1.0 | 0.98 | 0.99 | 1.0 | 1.0 | 1.0 |
| $f = 0.07$ | 1.0 | 1.0 | 1.0 | 1.0 | 1.0 | 1.0 |
| $f = 0.08$ | 1.0 | 1.0 | 1.0 | 1.0 | 1.0 | 1.0 |
| $f = 0.09$ | 1.0 | 1.0 | 1.0 | 1.0 | 1.0 | 1.0 |
| $f = 0.1$ | 1.0 | 1.0 | 1.0 | 1.0 | 1.0 | 1.0 |
| $f = 0.2$ | 1.0 | 1.0 | 1.0 | 1.0 | 1.0 | 1.0 |
| $f = 0.3$ | 1.0 | 1.0 | 1.0 | 1.0 | 1.0 | 1.0 |
| $f = 0.4$ | 1.0 | 1.0 | 1.0 | 1.0 | 1.0 | 1.0 |
| $f = 0.5$ | 1.0 | 1.0 | 1.0 | 1.0 | 1.0 | 1.0 |
| Original Model | 0.94 | 0.52 | 0.77 | 1.0 | 0.98 | 1.0 |

Table S8: Statistical power of $D$, $D_{anc}$, and $D+$ to detect introgression in the CEU population under realistic simulations of human demography (see Figure S2). The rows correspond to the simulated admixture proportions and the columns represent the proportion of statistically significant replicates from zero at *p-value*<0.05 out of 100 replicate simulations. Note that "Original Model" corresponds to the results using the original model from *Ragsdale and Gravel 2019* (see Figure S3).

| | $n = 1$ | | | $n = 100$ | | |
|---|---|---|---|---|---|---|
| | $D$ | $D_{anc}$ | $D+$ | $D$ | $D_{anc}$ | $D+$ |
| $f = 0$ | 0.16 | 0.07 | 0.11 | 0.0 | 0.0 | 0.0 |
| $f = 0.01$ | 0.49 | 0.26 | 0.36 | 0.69 | 0.18 | 0.47 |
| $f = 0.02$ | 0.91 | 0.54 | 0.76 | 0.96 | 0.5 | 0.83 |
| $f = 0.03$ | 0.97 | 0.77 | 0.87 | 1.0 | 0.9 | 0.98 |
| $f = 0.04$ | 1.0 | 0.86 | 0.95 | 1.0 | 0.93 | 0.98 |
| $f = 0.05$ | 1.0 | 0.91 | 0.97 | 1.0 | 0.99 | 1.0 |
| $f = 0.06$ | 1.0 | 0.97 | 0.99 | 1.0 | 1.0 | 1.0 |
| $f = 0.07$ | 1.0 | 1.0 | 1.0 | 1.0 | 1.0 | 1.0 |
| $f = 0.08$ | 1.0 | 1.0 | 1.0 | 1.0 | 1.0 | 1.0 |
| $f = 0.09$ | 1.0 | 1.0 | 1.0 | 1.0 | 1.0 | 1.0 |
| $f = 0.1$ | 1.0 | 1.0 | 1.0 | 1.0 | 1.0 | 1.0 |
| $f = 0.2$ | 1.0 | 1.0 | 1.0 | 1.0 | 1.0 | 1.0 |
| $f = 0.3$ | 1.0 | 1.0 | 1.0 | 1.0 | 1.0 | 1.0 |
| $f = 0.4$ | 1.0 | 1.0 | 1.0 | 1.0 | 1.0 | 1.0 |
| $f = 0.5$ | 1.0 | 1.0 | 1.0 | 1.0 | 1.0 | 1.0 |
| Original Model | 0.96 | 0.62 | 0.85 | 1.0 | 1.0 | 1.0 |

Table S9: Statistical power of $D$, $D_{anc}$, and $D+$ to detect introgression in the CHB population under realistic simulations of human demography (see Figure S2). The rows correspond to the simulated admixture proportions and the columns represent the proportion of statistically significant replicates from zero at *p-value*<0.05 out of 100 replicate simulations. Note that "Original Model" corresponds to the results using the original model from *Ragsdale and Gravel 2019* (see Figure S3).

| | $n = 1$ | | | | | | | | | $n = 100$ | | | | | | | | |
|---|---|---|---|---|---|---|---|---|---|---|---|---|---|---|---|---|---|---|
| | $D$ | | | $D_{anc}$ | | | $D+$ | | | $D$ | | | $D_{anc}$ | | | $D+$ | | |
| | $\mathbb{E}$ | $\mu$ | $\sigma$ | $\mathbb{E}$ | $\mu$ | $\sigma$ | $\mathbb{E}$ | $\mu$ | $\sigma$ | $\mathbb{E}$ | $\mu$ | $\sigma$ | $\mathbb{E}$ | $\mu$ | $\sigma$ | $\mathbb{E}$ | $\mu$ | $\sigma$ |
| $f = 0$ | 0.0 | -0.003 | 0.02 | 0.0 | -0.001 | 0.005 | 0.0 | -0.001 | 0.006 | 0.0 | 0.001 | 0.009 | 0.0 | 0.0 | 0.002 | 0.0 | 0.0 | 0.003 |
| $f = 0.01$ | 0.019 | 0.021 | 0.021 | 0.004 | 0.004 | 0.006 | 0.006 | 0.007 | 0.007 | 0.019 | 0.017 | 0.009 | 0.004 | 0.003 | 0.002 | 0.006 | 0.005 | 0.003 |
| $f = 0.02$ | 0.038 | 0.037 | 0.022 | 0.007 | 0.006 | 0.006 | 0.012 | 0.011 | 0.007 | 0.038 | 0.039 | 0.011 | 0.007 | 0.007 | 0.003 | 0.012 | 0.012 | 0.004 |
| $f = 0.03$ | 0.056 | 0.057 | 0.021 | 0.011 | 0.011 | 0.005 | 0.018 | 0.018 | 0.007 | 0.056 | 0.055 | 0.011 | 0.011 | 0.01 | 0.003 | 0.018 | 0.018 | 0.004 |
| $f = 0.04$ | 0.073 | 0.075 | 0.023 | 0.014 | 0.015 | 0.005 | 0.024 | 0.024 | 0.007 | 0.073 | 0.073 | 0.012 | 0.014 | 0.014 | 0.003 | 0.024 | 0.023 | 0.004 |
| $f = 0.05$ | 0.09 | 0.088 | 0.023 | 0.017 | 0.017 | 0.006 | 0.029 | 0.029 | 0.008 | 0.09 | 0.089 | 0.013 | 0.017 | 0.017 | 0.003 | 0.029 | 0.029 | 0.004 |
| $f = 0.06$ | 0.106 | 0.107 | 0.027 | 0.021 | 0.02 | 0.007 | 0.035 | 0.034 | 0.01 | 0.106 | 0.105 | 0.013 | 0.021 | 0.021 | 0.003 | 0.035 | 0.035 | 0.005 |
| $f = 0.07$ | 0.122 | 0.123 | 0.023 | 0.024 | 0.025 | 0.007 | 0.041 | 0.041 | 0.009 | 0.122 | 0.122 | 0.013 | 0.024 | 0.024 | 0.003 | 0.041 | 0.041 | 0.005 |
| $f = 0.08$ | 0.137 | 0.138 | 0.025 | 0.028 | 0.028 | 0.006 | 0.046 | 0.047 | 0.009 | 0.137 | 0.137 | 0.013 | 0.028 | 0.027 | 0.003 | 0.046 | 0.046 | 0.005 |
| $f = 0.09$ | 0.152 | 0.153 | 0.027 | 0.031 | 0.031 | 0.007 | 0.052 | 0.052 | 0.01 | 0.152 | 0.152 | 0.015 | 0.031 | 0.031 | 0.004 | 0.052 | 0.052 | 0.005 |
| $f = 0.1$ | 0.166 | 0.164 | 0.026 | 0.035 | 0.034 | 0.007 | 0.057 | 0.057 | 0.009 | 0.166 | 0.167 | 0.014 | 0.035 | 0.035 | 0.004 | 0.057 | 0.057 | 0.005 |
| $f = 0.2$ | 0.287 | 0.286 | 0.026 | 0.067 | 0.067 | 0.007 | 0.109 | 0.109 | 0.011 | 0.287 | 0.289 | 0.013 | 0.067 | 0.068 | 0.004 | 0.109 | 0.11 | 0.006 |
| $f = 0.3$ | 0.379 | 0.378 | 0.023 | 0.099 | 0.099 | 0.008 | 0.157 | 0.156 | 0.011 | 0.379 | 0.38 | 0.013 | 0.099 | 0.099 | 0.005 | 0.157 | 0.157 | 0.007 |
| $f = 0.4$ | 0.452 | 0.45 | 0.019 | 0.128 | 0.127 | 0.009 | 0.2 | 0.198 | 0.012 | 0.452 | 0.453 | 0.01 | 0.128 | 0.129 | 0.004 | 0.2 | 0.2 | 0.006 |
| $f = 0.5$ | 0.511 | 0.509 | 0.019 | 0.157 | 0.156 | 0.009 | 0.24 | 0.239 | 0.012 | 0.511 | 0.51 | 0.009 | 0.157 | 0.157 | 0.005 | 0.24 | 0.24 | 0.006 |

Table S10: Summary of methods to detect introgression in simulations under an IUA model of introgression (see Figure S1). The rows correspond to the simulated admixture proportions and the columns represent the coalescent based expectation ($\mathbb{E}$), mean ($\mu$), and standard deviation ($\sigma$) of metrics to detect introgression from 100 replicate simulations per admixture portion ($f$).

| | $n = 1$ | | | | | | $n = 100$ | | | | | |
|---|---|---|---|---|---|---|---|---|---|---|---|---|
| | $D$ | | $D_{anc}$ | | $D+$ | | $D$ | | $D_{anc}$ | | $D+$ | |
| | $\mu$ | $\sigma$ | $\mu$ | $\sigma$ | $\mu$ | $\sigma$ | $\mu$ | $\sigma$ | $\mu$ | $\sigma$ | $\mu$ | $\sigma$ |
| $f = 0$ | 0.005 | 0.041 | 0.0 | 0.006 | 0.001 | 0.006 | 0.0 | 0.014 | -0.0 | 0.002 | -0.0 | 0.002 |
| $f = 0.01$ | 0.089 | 0.071 | 0.005 | 0.006 | 0.009 | 0.008 | 0.101 | 0.055 | 0.005 | 0.004 | 0.01 | 0.007 |
| $f = 0.02$ | 0.173 | 0.09 | 0.009 | 0.008 | 0.018 | 0.012 | 0.18 | 0.059 | 0.01 | 0.005 | 0.019 | 0.009 |
| $f = 0.03$ | 0.24 | 0.076 | 0.015 | 0.008 | 0.028 | 0.013 | 0.247 | 0.052 | 0.015 | 0.005 | 0.029 | 0.009 |
| $f = 0.04$ | 0.3 | 0.079 | 0.021 | 0.009 | 0.039 | 0.014 | 0.298 | 0.055 | 0.02 | 0.006 | 0.038 | 0.011 |
| $f = 0.05$ | 0.339 | 0.074 | 0.025 | 0.01 | 0.047 | 0.016 | 0.347 | 0.049 | 0.026 | 0.006 | 0.048 | 0.011 |
| $f = 0.06$ | 0.371 | 0.078 | 0.029 | 0.01 | 0.055 | 0.016 | 0.376 | 0.046 | 0.029 | 0.006 | 0.054 | 0.011 |
| $f = 0.07$ | 0.402 | 0.07 | 0.034 | 0.011 | 0.062 | 0.017 | 0.411 | 0.041 | 0.035 | 0.007 | 0.064 | 0.012 |
| $f = 0.08$ | 0.443 | 0.061 | 0.039 | 0.01 | 0.073 | 0.016 | 0.433 | 0.043 | 0.038 | 0.007 | 0.07 | 0.013 |
| $f = 0.09$ | 0.457 | 0.069 | 0.043 | 0.013 | 0.079 | 0.021 | 0.46 | 0.045 | 0.043 | 0.009 | 0.08 | 0.015 |
| $f = 0.1$ | 0.487 | 0.067 | 0.049 | 0.011 | 0.089 | 0.019 | 0.485 | 0.041 | 0.048 | 0.008 | 0.087 | 0.014 |
| $f = 0.2$ | 0.623 | 0.051 | 0.093 | 0.015 | 0.161 | 0.024 | 0.617 | 0.028 | 0.091 | 0.012 | 0.159 | 0.019 |
| $f = 0.3$ | 0.684 | 0.038 | 0.132 | 0.015 | 0.222 | 0.022 | 0.686 | 0.019 | 0.133 | 0.012 | 0.222 | 0.017 |
| $f = 0.4$ | 0.726 | 0.039 | 0.169 | 0.017 | 0.275 | 0.025 | 0.728 | 0.015 | 0.17 | 0.01 | 0.275 | 0.014 |
| $f = 0.5$ | 0.766 | 0.03 | 0.206 | 0.017 | 0.325 | 0.022 | 0.758 | 0.013 | 0.203 | 0.01 | 0.32 | 0.013 |
| Original Model | 0.177 | 0.087 | 0.011 | 0.009 | 0.021 | 0.014 | 0.194 | 0.053 | 0.012 | 0.004 | 0.022 | 0.008 |

Table S11: Summary of methods to detect introgression in the CEU population under realistic simulations of human demography (see Figure S2). The rows correspond to the simulated admixture proportions and the columns represent the mean ($\mu$) and standard deviation ($\sigma$) of metrics to detect introgression from 100 replicate simulations per admixture portion ($f$). Note that "Original Model" corresponds to the results using the original model from *Ragsdale and Gravel 2019* (see Figure S3).

| | $n=1$ | | | | | | $n=100$ | | | | | |
|---|---|---|---|---|---|---|---|---|---|---|---|---|
| | $D$ | | $D_{anc}$ | | $D+$ | | $D$ | | $D_{anc}$ | | $D+$ | |
| | $\mu$ | $\sigma$ | $\mu$ | $\sigma$ | $\mu$ | $\sigma$ | $\mu$ | $\sigma$ | $\mu$ | $\sigma$ | $\mu$ | $\sigma$ |
| $f=0$ | 0.004 | 0.041 | -0.001 | 0.005 | -0.001 | 0.005 | -0.001 | 0.017 | -0.0 | 0.002 | -0.0 | 0.002 |
| $f=0.01$ | 0.09 | 0.072 | 0.005 | 0.006 | 0.009 | 0.008 | 0.101 | 0.057 | 0.005 | 0.004 | 0.01 | 0.006 |
| $f=0.02$ | 0.178 | 0.086 | 0.01 | 0.008 | 0.019 | 0.012 | 0.178 | 0.061 | 0.01 | 0.005 | 0.019 | 0.008 |
| $f=0.03$ | 0.247 | 0.094 | 0.016 | 0.01 | 0.03 | 0.015 | 0.251 | 0.059 | 0.016 | 0.006 | 0.03 | 0.01 |
| $f=0.04$ | 0.294 | 0.093 | 0.021 | 0.01 | 0.039 | 0.016 | 0.295 | 0.066 | 0.02 | 0.007 | 0.037 | 0.012 |
| $f=0.05$ | 0.344 | 0.086 | 0.026 | 0.011 | 0.048 | 0.018 | 0.346 | 0.055 | 0.026 | 0.006 | 0.048 | 0.011 |
| $f=0.06$ | 0.382 | 0.088 | 0.031 | 0.012 | 0.057 | 0.02 | 0.372 | 0.056 | 0.029 | 0.008 | 0.054 | 0.013 |
| $f=0.07$ | 0.405 | 0.081 | 0.035 | 0.012 | 0.063 | 0.019 | 0.41 | 0.054 | 0.035 | 0.008 | 0.064 | 0.014 |
| $f=0.08$ | 0.436 | 0.077 | 0.039 | 0.012 | 0.072 | 0.019 | 0.433 | 0.054 | 0.038 | 0.008 | 0.07 | 0.015 |
| $f=0.09$ | 0.465 | 0.071 | 0.044 | 0.013 | 0.081 | 0.021 | 0.461 | 0.049 | 0.044 | 0.009 | 0.08 | 0.015 |
| $f=0.1$ | 0.489 | 0.075 | 0.05 | 0.013 | 0.09 | 0.023 | 0.488 | 0.05 | 0.049 | 0.01 | 0.088 | 0.017 |
| $f=0.2$ | 0.625 | 0.052 | 0.094 | 0.015 | 0.163 | 0.023 | 0.624 | 0.031 | 0.094 | 0.013 | 0.163 | 0.02 |
| $f=0.3$ | 0.688 | 0.042 | 0.134 | 0.018 | 0.224 | 0.027 | 0.688 | 0.025 | 0.135 | 0.014 | 0.225 | 0.02 |
| $f=0.4$ | 0.733 | 0.036 | 0.173 | 0.018 | 0.28 | 0.025 | 0.732 | 0.019 | 0.173 | 0.013 | 0.28 | 0.019 |
| $f=0.5$ | 0.771 | 0.033 | 0.211 | 0.019 | 0.331 | 0.026 | 0.762 | 0.017 | 0.206 | 0.015 | 0.324 | 0.019 |
| Original Model | 0.189 | 0.081 | 0.012 | 0.008 | 0.023 | 0.014 | 0.199 | 0.052 | 0.012 | 0.005 | 0.023 | 0.008 |

Table S12: Summary of methods to detect introgression in the CHB population under realistic simulations of human demography (see Figure S2). The rows correspond to the simulated admixture proportions and the columns represent the mean ($\mu$) and standard deviation ($\sigma$) of metrics to detect introgression from 100 replicate simulations per admixture portion ($f$). Note that "Original Model" corresponds to the results using the original model from *Ragsdale and Gravel 2019* (see Figure S3).

| | $n = 1$ | | | | | | $n = 100$ | | | | | |
|---|---|---|---|---|---|---|---|---|---|---|---|---|
| | $f_{hom}$ | | $f_{anc}$ | | $f+$ | | $f_{hom}$ | | $f_{anc}$ | | $f+$ | |
| | $RMSE$ | $MAE$ | $RMSE$ | $MAE$ | $RMSE$ | $MAE$ | $RMSE$ | $MAE$ | $RMSE$ | $MAE$ | $RMSE$ | $MAE$ |
| $f = 0$ | 0.004 | 0.003 | 0.006 | 0.005 | 0.004 | 0.003 | 0.002 | 0.001 | 0.002 | 0.002 | 0.002 | 0.001 |
| $f = 0.01$ | 0.007 | 0.006 | 0.008 | 0.007 | 0.007 | 0.006 | 0.007 | 0.006 | 0.007 | 0.006 | 0.007 | 0.006 |
| $f = 0.02$ | 0.013 | 0.012 | 0.014 | 0.013 | 0.013 | 0.012 | 0.012 | 0.012 | 0.013 | 0.012 | 0.012 | 0.012 |
| $f = 0.03$ | 0.018 | 0.018 | 0.019 | 0.018 | 0.018 | 0.018 | 0.018 | 0.018 | 0.018 | 0.018 | 0.018 | 0.018 |
| $f = 0.04$ | 0.024 | 0.024 | 0.024 | 0.023 | 0.024 | 0.023 | 0.024 | 0.024 | 0.024 | 0.024 | 0.024 | 0.024 |
| $f = 0.05$ | 0.031 | 0.03 | 0.031 | 0.03 | 0.031 | 0.03 | 0.03 | 0.03 | 0.031 | 0.03 | 0.031 | 0.03 |
| $f = 0.06$ | 0.036 | 0.036 | 0.038 | 0.037 | 0.037 | 0.036 | 0.037 | 0.036 | 0.036 | 0.036 | 0.036 | 0.036 |
| $f = 0.07$ | 0.042 | 0.042 | 0.042 | 0.042 | 0.042 | 0.042 | 0.042 | 0.042 | 0.042 | 0.042 | 0.042 | 0.042 |
| $f = 0.08$ | 0.048 | 0.048 | 0.048 | 0.048 | 0.048 | 0.048 | 0.048 | 0.048 | 0.049 | 0.048 | 0.048 | 0.048 |
| $f = 0.09$ | 0.054 | 0.054 | 0.055 | 0.054 | 0.054 | 0.054 | 0.054 | 0.054 | 0.054 | 0.054 | 0.054 | 0.054 |
| $f = 0.1$ | 0.061 | 0.06 | 0.061 | 0.06 | 0.061 | 0.06 | 0.06 | 0.06 | 0.06 | 0.06 | 0.06 | 0.06 |
| $f = 0.2$ | 0.12 | 0.12 | 0.12 | 0.12 | 0.12 | 0.12 | 0.119 | 0.119 | 0.12 | 0.12 | 0.12 | 0.119 |
| $f = 0.3$ | 0.181 | 0.18 | 0.18 | 0.18 | 0.18 | 0.18 | 0.18 | 0.179 | 0.179 | 0.179 | 0.179 | 0.179 |
| $f = 0.4$ | 0.241 | 0.241 | 0.242 | 0.241 | 0.241 | 0.241 | 0.24 | 0.24 | 0.24 | 0.24 | 0.24 | 0.24 |
| $f = 0.5$ | 0.302 | 0.301 | 0.301 | 0.301 | 0.301 | 0.301 | 0.3 | 0.3 | 0.3 | 0.3 | 0.3 | 0.3 |

Table S13: Statistical power of $f_{hom}$, $f_{anc}$, and $f+$ to infer the true admixture proportion in simulations under an IUA model of introgression (see Figure S1). The rows correspond to the simulated admixture proportions and the columns represent either the root-mean-squared-error ($RMSE$) or mean-absolute-error ($MAE$) for each statistic.

| | $n = 1$ | | | | | | $n = 100$ | | | | | |
|---|---|---|---|---|---|---|---|---|---|---|---|---|
| | $f_{hom}$ | | $f_{anc}$ | | $f+$ | | $f_{hom}$ | | $f_{anc}$ | | $f+$ | |
| | *RMSE* | *MAE* | *RMSE* | *MAE* | *RMSE* | *MAE* | *RMSE* | *MAE* | *RMSE* | *MAE* | *RMSE* | *MAE* |
| $f = 0$ | 0.002 | 0.002 | 0.006 | 0.005 | 0.003 | 0.003 | 0.001 | 0.001 | 0.002 | 0.002 | 0.001 | 0.001 |
| $f = 0.01$ | 0.007 | 0.006 | 0.008 | 0.006 | 0.007 | 0.006 | 0.006 | 0.005 | 0.006 | 0.005 | 0.006 | 0.005 |
| $f = 0.02$ | 0.012 | 0.01 | 0.013 | 0.011 | 0.012 | 0.011 | 0.01 | 0.01 | 0.01 | 0.01 | 0.01 | 0.01 |
| $f = 0.03$ | 0.016 | 0.014 | 0.016 | 0.014 | 0.016 | 0.014 | 0.014 | 0.013 | 0.015 | 0.013 | 0.014 | 0.013 |
| $f = 0.04$ | 0.02 | 0.018 | 0.02 | 0.018 | 0.02 | 0.018 | 0.019 | 0.018 | 0.019 | 0.018 | 0.019 | 0.018 |
| $f = 0.05$ | 0.025 | 0.023 | 0.025 | 0.023 | 0.025 | 0.023 | 0.023 | 0.022 | 0.023 | 0.022 | 0.023 | 0.022 |
| $f = 0.06$ | 0.03 | 0.028 | 0.03 | 0.028 | 0.03 | 0.028 | 0.029 | 0.028 | 0.029 | 0.028 | 0.029 | 0.028 |
| $f = 0.07$ | 0.035 | 0.034 | 0.036 | 0.033 | 0.035 | 0.034 | 0.033 | 0.032 | 0.033 | 0.032 | 0.033 | 0.032 |
| $f = 0.08$ | 0.038 | 0.036 | 0.039 | 0.037 | 0.038 | 0.037 | 0.039 | 0.039 | 0.04 | 0.039 | 0.039 | 0.039 |
| $f = 0.09$ | 0.044 | 0.042 | 0.045 | 0.043 | 0.045 | 0.043 | 0.043 | 0.042 | 0.044 | 0.042 | 0.043 | 0.042 |
| $f = 0.1$ | 0.048 | 0.046 | 0.048 | 0.047 | 0.048 | 0.047 | 0.048 | 0.047 | 0.048 | 0.047 | 0.048 | 0.047 |
| $f = 0.2$ | 0.097 | 0.095 | 0.097 | 0.096 | 0.097 | 0.095 | 0.099 | 0.098 | 0.098 | 0.097 | 0.098 | 0.097 |
| $f = 0.3$ | 0.149 | 0.148 | 0.149 | 0.148 | 0.149 | 0.148 | 0.148 | 0.147 | 0.148 | 0.147 | 0.148 | 0.147 |
| $f = 0.4$ | 0.202 | 0.201 | 0.203 | 0.202 | 0.203 | 0.201 | 0.202 | 0.201 | 0.202 | 0.201 | 0.202 | 0.201 |
| $f = 0.5$ | 0.255 | 0.254 | 0.255 | 0.254 | 0.255 | 0.254 | 0.259 | 0.258 | 0.258 | 0.258 | 0.259 | 0.258 |
| Original Model | 0.009 | 0.007 | 0.011 | 0.008 | 0.01 | 0.007 | 0.008 | 0.007 | 0.008 | 0.007 | 0.008 | 0.007 |

Table S14: Statistical power of $f_{hom}$, $f_{anc}$, and $f+$ to infer the true admixture proportion in the CEU population under realistic simulations of human demography (see Figure S2). The rows correspond to the simulated admixture proportions and the columns represent either the root-mean-squared-error (*RMSE*) or mean-absolute-error (*MAE*) for each statistic. Note that "Original Model" corresponds to the results using the original model from *Ragsdale and Gravel 2019* (see Figure S3).

| | $n=1$ | | | | | | $n=100$ | | | | | |
|---|---|---|---|---|---|---|---|---|---|---|---|---|
| | $f_{hom}$ | | $f_{anc}$ | | $f+$ | | $f_{hom}$ | | $f_{anc}$ | | $f+$ | |
| | *RMSE* | *MAE* | *RMSE* | *MAE* | *RMSE* | *MAE* | *RMSE* | *MAE* | *RMSE* | *MAE* | *RMSE* | *MAE* |
| $f=0$ | 0.002 | 0.002 | 0.005 | 0.004 | 0.003 | 0.002 | 0.001 | 0.001 | 0.002 | 0.002 | 0.001 | 0.001 |
| $f=0.01$ | 0.007 | 0.006 | 0.008 | 0.006 | 0.007 | 0.006 | 0.006 | 0.005 | 0.006 | 0.005 | 0.006 | 0.005 |
| $f=0.02$ | 0.011 | 0.01 | 0.012 | 0.01 | 0.011 | 0.01 | 0.01 | 0.009 | 0.01 | 0.009 | 0.01 | 0.009 |
| $f=0.03$ | 0.015 | 0.014 | 0.016 | 0.013 | 0.015 | 0.013 | 0.014 | 0.013 | 0.014 | 0.013 | 0.014 | 0.013 |
| $f=0.04$ | 0.02 | 0.018 | 0.021 | 0.018 | 0.02 | 0.018 | 0.02 | 0.018 | 0.02 | 0.019 | 0.02 | 0.019 |
| $f=0.05$ | 0.024 | 0.022 | 0.025 | 0.023 | 0.025 | 0.022 | 0.023 | 0.022 | 0.023 | 0.022 | 0.023 | 0.022 |
| $f=0.06$ | 0.029 | 0.027 | 0.03 | 0.027 | 0.029 | 0.027 | 0.03 | 0.028 | 0.03 | 0.029 | 0.03 | 0.028 |
| $f=0.07$ | 0.035 | 0.033 | 0.035 | 0.032 | 0.035 | 0.033 | 0.033 | 0.032 | 0.033 | 0.032 | 0.033 | 0.032 |
| $f=0.08$ | 0.039 | 0.037 | 0.04 | 0.037 | 0.039 | 0.037 | 0.039 | 0.038 | 0.039 | 0.038 | 0.039 | 0.038 |
| $f=0.09$ | 0.043 | 0.041 | 0.044 | 0.042 | 0.044 | 0.041 | 0.043 | 0.042 | 0.043 | 0.042 | 0.043 | 0.042 |
| $f=0.1$ | 0.048 | 0.046 | 0.048 | 0.046 | 0.048 | 0.046 | 0.048 | 0.047 | 0.048 | 0.047 | 0.048 | 0.047 |
| $f=0.2$ | 0.096 | 0.094 | 0.096 | 0.094 | 0.096 | 0.094 | 0.096 | 0.095 | 0.095 | 0.094 | 0.096 | 0.094 |
| $f=0.3$ | 0.147 | 0.146 | 0.147 | 0.146 | 0.147 | 0.146 | 0.146 | 0.145 | 0.146 | 0.145 | 0.146 | 0.145 |
| $f=0.4$ | 0.198 | 0.197 | 0.198 | 0.197 | 0.198 | 0.197 | 0.198 | 0.197 | 0.198 | 0.197 | 0.198 | 0.197 |
| $f=0.5$ | 0.25 | 0.249 | 0.249 | 0.248 | 0.249 | 0.248 | 0.255 | 0.254 | 0.254 | 0.254 | 0.255 | 0.254 |
| Original Model | 0.01 | 0.007 | 0.012 | 0.009 | 0.011 | 0.008 | 0.009 | 0.007 | 0.009 | 0.007 | 0.009 | 0.007 |

Table S15: Statistical power of $f_{hom}$, $f_{anc}$, and $f+$ to infer the true admixture proportion in the CHB population under realistic simulations of human demography (see Figure S2). The rows correspond to the simulated admixture proportions and the columns represent either the root-mean-squared-error (*RMSE*) or mean-absolute-error (*MAE*) for each statistic. Note that "Original Model" corresponds to the results using the original model from *Ragsdale and Gravel 2019* (see Figure S3).

| | $n = 1$ | | | | | | | | | $n = 100$ | | | | | | | | |
|---|---|---|---|---|---|---|---|---|---|---|---|---|---|---|---|---|---|---|
| | $f_{hom}$ | | | $f_{anc}$ | | | $f+$ | | | $f_{hom}$ | | | $f_{anc}$ | | | $f+$ | | |
| | $\mathbb{E}$ | $\mu$ | $\sigma$ | $\mathbb{E}$ | $\mu$ | $\sigma$ | $\mathbb{E}$ | $\mu$ | $\sigma$ | $\mathbb{E}$ | $\mu$ | $\sigma$ | $\mathbb{E}$ | $\mu$ | $\sigma$ | $\mathbb{E}$ | $\mu$ | $\sigma$ |
| $f = 0$ | 0.0 | -0.001 | 0.004 | 0.0 | -0.001 | 0.005 | 0.0 | -0.001 | 0.004 | 0.0 | 0.0 | 0.002 | 0.0 | 0.0 | 0.002 | 0.0 | 0.0 | 0.002 |
| $f = 0.01$ | 0.004 | 0.004 | 0.004 | 0.004 | 0.005 | 0.006 | 0.004 | 0.004 | 0.005 | 0.004 | 0.004 | 0.002 | 0.004 | 0.004 | 0.003 | 0.004 | 0.004 | 0.002 |
| $f = 0.02$ | 0.008 | 0.008 | 0.005 | 0.008 | 0.007 | 0.007 | 0.008 | 0.008 | 0.005 | 0.008 | 0.008 | 0.002 | 0.008 | 0.008 | 0.003 | 0.008 | 0.008 | 0.002 |
| $f = 0.03$ | 0.012 | 0.012 | 0.004 | 0.012 | 0.012 | 0.006 | 0.012 | 0.012 | 0.005 | 0.012 | 0.012 | 0.002 | 0.012 | 0.012 | 0.003 | 0.012 | 0.012 | 0.003 |
| $f = 0.04$ | 0.016 | 0.016 | 0.005 | 0.016 | 0.017 | 0.006 | 0.016 | 0.017 | 0.005 | 0.016 | 0.016 | 0.003 | 0.016 | 0.016 | 0.003 | 0.016 | 0.016 | 0.003 |
| $f = 0.05$ | 0.02 | 0.02 | 0.005 | 0.02 | 0.02 | 0.006 | 0.02 | 0.02 | 0.005 | 0.02 | 0.02 | 0.003 | 0.02 | 0.02 | 0.004 | 0.02 | 0.02 | 0.003 |
| $f = 0.06$ | 0.024 | 0.024 | 0.006 | 0.024 | 0.023 | 0.008 | 0.024 | 0.024 | 0.007 | 0.024 | 0.024 | 0.003 | 0.024 | 0.024 | 0.004 | 0.024 | 0.024 | 0.003 |
| $f = 0.07$ | 0.028 | 0.028 | 0.006 | 0.028 | 0.028 | 0.008 | 0.028 | 0.028 | 0.006 | 0.028 | 0.028 | 0.003 | 0.028 | 0.028 | 0.004 | 0.028 | 0.028 | 0.003 |
| $f = 0.08$ | 0.032 | 0.032 | 0.006 | 0.032 | 0.032 | 0.007 | 0.032 | 0.032 | 0.006 | 0.032 | 0.032 | 0.003 | 0.032 | 0.032 | 0.004 | 0.032 | 0.032 | 0.004 |
| $f = 0.09$ | 0.036 | 0.036 | 0.007 | 0.036 | 0.036 | 0.008 | 0.036 | 0.036 | 0.007 | 0.036 | 0.036 | 0.004 | 0.036 | 0.036 | 0.004 | 0.036 | 0.036 | 0.004 |
| $f = 0.1$ | 0.04 | 0.04 | 0.007 | 0.04 | 0.04 | 0.008 | 0.04 | 0.04 | 0.007 | 0.04 | 0.04 | 0.004 | 0.04 | 0.04 | 0.004 | 0.04 | 0.04 | 0.004 |
| $f = 0.2$ | 0.08 | 0.08 | 0.008 | 0.08 | 0.08 | 0.009 | 0.08 | 0.08 | 0.008 | 0.08 | 0.081 | 0.005 | 0.08 | 0.08 | 0.005 | 0.08 | 0.081 | 0.005 |
| $f = 0.3$ | 0.12 | 0.12 | 0.009 | 0.12 | 0.12 | 0.01 | 0.12 | 0.12 | 0.009 | 0.12 | 0.121 | 0.006 | 0.12 | 0.121 | 0.006 | 0.12 | 0.121 | 0.006 |
| $f = 0.4$ | 0.16 | 0.159 | 0.01 | 0.16 | 0.159 | 0.011 | 0.16 | 0.159 | 0.01 | 0.16 | 0.16 | 0.005 | 0.16 | 0.16 | 0.006 | 0.16 | 0.16 | 0.005 |
| $f = 0.5$ | 0.2 | 0.199 | 0.012 | 0.2 | 0.199 | 0.012 | 0.2 | 0.199 | 0.011 | 0.2 | 0.2 | 0.006 | 0.2 | 0.2 | 0.006 | 0.2 | 0.2 | 0.006 |

Table S16: Summary of methods to quantify introgression in simulations under an IUA model of introgression (see Figure S1). The rows correspond to the simulated admixture proportions and the columns represent the coalescent based expectation ($\mathbb{E}$), mean ($\mu$), and standard deviation ($\sigma$) of metrics to quantify introgression from 100 replicate simulations per admixture portion ($f$).

| | $n = 1$ | | | | | | $n = 100$ | | | | | |
|---|---|---|---|---|---|---|---|---|---|---|---|---|
| | $f_{hom}$ | | $f_{anc}$ | | $f+$ | | $f_{hom}$ | | $f_{anc}$ | | $f+$ | |
| | $\mu$ | $\sigma$ | $\mu$ | $\sigma$ | $\mu$ | $\sigma$ | $\mu$ | $\sigma$ | $\mu$ | $\sigma$ | $\mu$ | $\sigma$ |
| $f = 0$ | 0.0 | 0.002 | 0.0 | 0.006 | 0.0 | 0.003 | 0.0 | 0.001 | -0.0 | 0.002 | -0.0 | 0.001 |
| $f = 0.01$ | 0.005 | 0.004 | 0.006 | 0.006 | 0.005 | 0.005 | 0.006 | 0.003 | 0.005 | 0.004 | 0.005 | 0.004 |
| $f = 0.02$ | 0.011 | 0.007 | 0.01 | 0.008 | 0.01 | 0.007 | 0.011 | 0.005 | 0.011 | 0.005 | 0.011 | 0.005 |
| $f = 0.03$ | 0.016 | 0.007 | 0.016 | 0.009 | 0.016 | 0.007 | 0.017 | 0.005 | 0.017 | 0.006 | 0.017 | 0.005 |
| $f = 0.04$ | 0.022 | 0.008 | 0.022 | 0.009 | 0.022 | 0.009 | 0.022 | 0.006 | 0.022 | 0.006 | 0.022 | 0.006 |
| $f = 0.05$ | 0.027 | 0.01 | 0.027 | 0.011 | 0.027 | 0.01 | 0.028 | 0.007 | 0.028 | 0.007 | 0.028 | 0.007 |
| $f = 0.06$ | 0.032 | 0.01 | 0.032 | 0.011 | 0.032 | 0.01 | 0.032 | 0.007 | 0.032 | 0.007 | 0.032 | 0.006 |
| $f = 0.07$ | 0.036 | 0.01 | 0.037 | 0.012 | 0.036 | 0.011 | 0.038 | 0.008 | 0.038 | 0.008 | 0.038 | 0.008 |
| $f = 0.08$ | 0.044 | 0.01 | 0.043 | 0.011 | 0.043 | 0.01 | 0.041 | 0.008 | 0.041 | 0.008 | 0.041 | 0.008 |
| $f = 0.09$ | 0.048 | 0.013 | 0.047 | 0.014 | 0.047 | 0.013 | 0.048 | 0.01 | 0.048 | 0.01 | 0.048 | 0.01 |
| $f = 0.1$ | 0.054 | 0.012 | 0.053 | 0.013 | 0.053 | 0.012 | 0.053 | 0.009 | 0.053 | 0.01 | 0.053 | 0.009 |
| $f = 0.2$ | 0.105 | 0.017 | 0.104 | 0.018 | 0.105 | 0.018 | 0.102 | 0.014 | 0.103 | 0.014 | 0.103 | 0.014 |
| $f = 0.3$ | 0.152 | 0.017 | 0.152 | 0.018 | 0.152 | 0.017 | 0.153 | 0.014 | 0.153 | 0.014 | 0.153 | 0.014 |
| $f = 0.4$ | 0.199 | 0.021 | 0.198 | 0.021 | 0.199 | 0.021 | 0.199 | 0.013 | 0.199 | 0.013 | 0.199 | 0.013 |
| $f = 0.5$ | 0.246 | 0.019 | 0.246 | 0.02 | 0.246 | 0.019 | 0.242 | 0.013 | 0.242 | 0.013 | 0.242 | 0.013 |
| Original Model | 0.012 | 0.007 | 0.012 | 0.009 | 0.012 | 0.008 | 0.013 | 0.005 | 0.013 | 0.005 | 0.013 | 0.005 |

Table S17: Summary of methods to quantify introgression in the CEU population under realistic simulations of human demography (see Figure S2). The rows correspond to the simulated admixture proportions and the columns represent the mean ($\mu$) and standard deviation ($\sigma$) of metrics to quantify introgression from 100 replicate simulations per admixture portion ($f$). Note that "Original Model" corresponds to the results using the original model from *Ragsdale and Gravel 2019* (see Figure S3).

| | $n = 1$ | | | | | | $n = 100$ | | | | | |
|---|---|---|---|---|---|---|---|---|---|---|---|---|
| | $f_{hom}$ | | $f_{anc}$ | | $f+$ | | $f_{hom}$ | | $f_{anc}$ | | $f+$ | |
| | $\mu$ | $\sigma$ | $\mu$ | $\sigma$ | $\mu$ | $\sigma$ | $\mu$ | $\sigma$ | $\mu$ | $\sigma$ | $\mu$ | $\sigma$ |
| $f = 0$ | 0.0 | 0.002 | -0.001 | 0.005 | -0.0 | 0.003 | -0.0 | 0.001 | -0.0 | 0.002 | -0.0 | 0.001 |
| $f = 0.01$ | 0.005 | 0.004 | 0.005 | 0.006 | 0.005 | 0.005 | 0.006 | 0.003 | 0.005 | 0.004 | 0.006 | 0.004 |
| $f = 0.02$ | 0.011 | 0.006 | 0.011 | 0.008 | 0.011 | 0.007 | 0.011 | 0.005 | 0.011 | 0.005 | 0.011 | 0.005 |
| $f = 0.03$ | 0.017 | 0.008 | 0.018 | 0.01 | 0.017 | 0.009 | 0.017 | 0.006 | 0.017 | 0.006 | 0.017 | 0.006 |
| $f = 0.04$ | 0.022 | 0.009 | 0.022 | 0.011 | 0.022 | 0.01 | 0.022 | 0.007 | 0.021 | 0.007 | 0.022 | 0.007 |
| $f = 0.05$ | 0.028 | 0.011 | 0.028 | 0.012 | 0.028 | 0.011 | 0.028 | 0.007 | 0.028 | 0.007 | 0.028 | 0.007 |
| $f = 0.06$ | 0.033 | 0.011 | 0.033 | 0.013 | 0.033 | 0.012 | 0.032 | 0.008 | 0.031 | 0.008 | 0.032 | 0.008 |
| $f = 0.07$ | 0.037 | 0.012 | 0.038 | 0.013 | 0.037 | 0.012 | 0.038 | 0.009 | 0.038 | 0.009 | 0.038 | 0.009 |
| $f = 0.08$ | 0.043 | 0.012 | 0.043 | 0.013 | 0.043 | 0.012 | 0.042 | 0.009 | 0.042 | 0.009 | 0.042 | 0.009 |
| $f = 0.09$ | 0.049 | 0.013 | 0.048 | 0.014 | 0.049 | 0.014 | 0.048 | 0.01 | 0.048 | 0.01 | 0.048 | 0.01 |
| $f = 0.1$ | 0.054 | 0.015 | 0.054 | 0.015 | 0.054 | 0.015 | 0.053 | 0.011 | 0.053 | 0.011 | 0.053 | 0.011 |
| $f = 0.2$ | 0.106 | 0.017 | 0.106 | 0.017 | 0.106 | 0.017 | 0.105 | 0.015 | 0.106 | 0.015 | 0.106 | 0.015 |
| $f = 0.3$ | 0.154 | 0.021 | 0.154 | 0.022 | 0.154 | 0.021 | 0.155 | 0.017 | 0.155 | 0.017 | 0.155 | 0.017 |
| $f = 0.4$ | 0.203 | 0.021 | 0.203 | 0.022 | 0.203 | 0.021 | 0.203 | 0.016 | 0.203 | 0.017 | 0.203 | 0.016 |
| $f = 0.5$ | 0.251 | 0.024 | 0.252 | 0.024 | 0.252 | 0.024 | 0.246 | 0.019 | 0.246 | 0.019 | 0.246 | 0.019 |
| Original Model | 0.013 | 0.008 | 0.013 | 0.009 | 0.013 | 0.008 | 0.013 | 0.005 | 0.013 | 0.005 | 0.013 | 0.005 |

Table S18: Summary of methods to quantify introgression in the CHB population under realistic simulations of human demography (see Figure S2). The rows correspond to the simulated admixture proportions and the columns represent the mean ($\mu$) and standard deviation ($\sigma$) of metrics to quantify introgression from 100 replicate simulations per admixture portion ($f$). Note that "Original Model" corresponds to the results using the original model from *Ragsdale and Gravel 2019* (see Figure S3).

| | EUR | | | | | | ASN | | | | | |
|---|---|---|---|---|---|---|---|---|---|---|---|---|
| | $n = 1$ | | | $n = 100$ | | | $n = 1$ | | | $n = 100$ | | |
| | $ABBA - BABA$ | $BAAA - ABAA$ | $p - value$ | $ABBA - BABA$ | $BAAA - ABAA$ | $p - value$ | $ABBA - BABA$ | $BAAA - ABAA$ | $p - value$ | $ABBA - BABA$ | $BAAA - ABAA$ | $p - value$ |
| $f_{NEA} = 0;\ f_{DEN} = 0$ | 1.13 | -3.74 | 0.493 | -2.148 | 0.981 | 0.49 | -23.1 | -5.79 | 0.475 | -5.75 | -13.134 | 0.474 |
| $f_{NEA} = 0;\ f_{DEN} = 0.005$ | 7.63 | -3.41 | 0.483 | 3.397 | 16.614 | 0.452 | 76.81 | 72.92 | 0.495 | 71.184 | 83.656 | 0.455 |
| $f_{NEA} = 0;\ f_{DEN} = 0.01$ | 10.27 | 25.77 | 0.477 | -4.663 | -16.074 | 0.463 | 134.88 | 139.63 | 0.493 | 126.192 | 116.47 | 0.468 |
| $f_{NEA} = 0;\ f_{DEN} = 0.015$ | 8.53 | -26.29 | 0.453 | -6.341 | -5.129 | 0.496 | 208.21 | 232.71 | 0.461 | 198.766 | 211.171 | 0.46 |
| $f_{NEA} = 0;\ f_{DEN} = 0.02$ | 23.45 | 40.21 | 0.473 | -1.392 | -8.143 | 0.476 | 310.77 | 291.55 | 0.474 | 272.539 | 262.174 | 0.463 |
| $f_{NEA} = 0.005;\ f_{DEN} = 0$ | 177.68 | 154.06 | 0.468 | 174.963 | 184.513 | 0.469 | 401.72 | 355.71 | 0.432 | 358.169 | 369.07 | 0.468 |
| $f_{NEA} = 0.005;\ f_{DEN} = 0.005$ | 151.68 | 151.42 | 0.5 | 183.871 | 181.543 | 0.493 | 420.86 | 423.17 | 0.497 | 438.172 | 438.623 | 0.499 |
| $f_{NEA} = 0.005;\ f_{DEN} = 0.01$ | 181.47 | 147.95 | 0.452 | 171.577 | 155.368 | 0.445 | 512.44 | 488.88 | 0.465 | 485.462 | 477.153 | 0.471 |
| $f_{NEA} = 0.005;\ f_{DEN} = 0.015$ | 194.65 | 183.37 | 0.482 | 187.383 | 190.038 | 0.491 | 588.01 | 616.88 | 0.456 | 574.423 | 575.571 | 0.496 |
| $f_{NEA} = 0.005;\ f_{DEN} = 0.02$ | 151.4 | 177.47 | 0.458 | 170.253 | 157.222 | 0.463 | 598.77 | 565.56 | 0.449 | 615.728 | 614.482 | 0.496 |
| $f_{NEA} = 0.01;\ f_{DEN} = 0$ | 348.61 | 295.65 | 0.425 | 361.37 | 360.53 | 0.497 | 708.15 | 706.5 | 0.498 | 729.426 | 724.445 | 0.484 |
| $f_{NEA} = 0.01;\ f_{DEN} = 0.005$ | 333.47 | 320.52 | 0.479 | 358.112 | 358.932 | 0.498 | 805.86 | 782.3 | 0.46 | 803.604 | 781.849 | 0.432 |
| $f_{NEA} = 0.01;\ f_{DEN} = 0.01$ | 345.26 | 360.64 | 0.475 | 375.792 | 386.991 | 0.465 | 873.63 | 872.38 | 0.498 | 868.126 | 876.717 | 0.474 |
| $f_{NEA} = 0.01;\ f_{DEN} = 0.015$ | 365.14 | 340.59 | 0.461 | 359.884 | 375.163 | 0.442 | 925.75 | 899.36 | 0.458 | 918.066 | 939.6 | 0.424 |
| $f_{NEA} = 0.01;\ f_{DEN} = 0.02$ | 352.47 | 386.22 | 0.45 | 362.348 | 350.404 | 0.466 | 993.27 | 995.7 | 0.497 | 999.34 | 985.154 | 0.453 |
| $f_{NEA} = 0.015;\ f_{DEN} = 0$ | 525.7 | 511.52 | 0.478 | 509.951 | 513.517 | 0.489 | 1106.77 | 1108.42 | 0.498 | 1066.485 | 1066.743 | 0.499 |
| $f_{NEA} = 0.015;\ f_{DEN} = 0.005$ | 570.63 | 525.48 | 0.432 | 559.119 | 570.418 | 0.461 | 1180.67 | 1179.24 | 0.498 | 1165.936 | 1179.591 | 0.446 |
| $f_{NEA} = 0.015;\ f_{DEN} = 0.01$ | 541.25 | 525.89 | 0.477 | 534.657 | 540.419 | 0.479 | 1213.38 | 1225.37 | 0.482 | 1202.914 | 1194.777 | 0.472 |
| $f_{NEA} = 0.015;\ f_{DEN} = 0.015$ | 532.38 | 525.03 | 0.487 | 550.304 | 548.61 | 0.495 | 1304.08 | 1325.41 | 0.467 | 1276.638 | 1267.785 | 0.471 |
| $f_{NEA} = 0.015;\ f_{DEN} = 0.02$ | 493.97 | 506.58 | 0.481 | 535.464 | 539.921 | 0.486 | 1348.94 | 1378.87 | 0.454 | 1325.467 | 1333.965 | 0.47 |
| $f_{NEA} = 0.02;\ f_{DEN} = 0$ | 695.03 | 684.67 | 0.483 | 718.255 | 716.803 | 0.495 | 1448.84 | 1422.47 | 0.46 | 1447.819 | 1442.411 | 0.483 |
| $f_{NEA} = 0.02;\ f_{DEN} = 0.005$ | 717.56 | 704.26 | 0.48 | 713.221 | 702.212 | 0.461 | 1536.15 | 1509.74 | 0.461 | 1507.578 | 1510.627 | 0.49 |
| $f_{NEA} = 0.02;\ f_{DEN} = 0.01$ | 752.92 | 768.5 | 0.475 | 749.628 | 756.473 | 0.474 | 1615.03 | 1600.18 | 0.48 | 1611.919 | 1629.797 | 0.433 |
| $f_{NEA} = 0.02;\ f_{DEN} = 0.015$ | 694.22 | 708.01 | 0.481 | 729.494 | 716.208 | 0.457 | 1616.21 | 1655.6 | 0.439 | 1647.817 | 1634.883 | 0.458 |
| $f_{NEA} = 0.02;\ f_{DEN} = 0.02$ | 735.42 | 729.57 | 0.493 | 742.87 | 762.032 | 0.437 | 1806.93 | 1803.72 | 0.495 | 1734.26 | 1745.281 | 0.464 |

Table S19: Assessing if *(ABBA − BABA)* − *(BAAA − ABAA)* significantly differs from zero in the hypothetical European and Asian populations under a multi-pulse model of introgression from Neanderthals and Denisovans (see Figure S4) for all pairwise combinations of admixture proportions ($f_{NEA}$ and $f_{DEN}$) from 0% to 2%. The rows correspond to the simulated admixture proportions, the columns represent the mean site pattern difference across 100 replicate simulations and the corresponding *p-value* to assess if *(ABBA − BABA)* − *(BAAA − ABAA)* significantly differs from zero. After accounting for multiple-comparisons using the Bonferroni correction, a *p-value* less than 0.002 is considered significant.

| | EUR | | | | | | ASN | | | | | |
|---|---|---|---|---|---|---|---|---|---|---|---|---|
| | $n = 1$ | | | $n = 100$ | | | $n = 1$ | | | $n = 100$ | | |
| | $ABBA - BABA$ | $BAAA - ABAA$ | $p - value$ | $ABBA - BABA$ | $BAAA - ABAA$ | $p - value$ | $ABBA - BABA$ | $BAAA - ABAA$ | $p - value$ | $ABBA - BABA$ | $BAAA - ABAA$ | $p - value$ |
| $f_{NEA} = 0;\ f_{DEN} = 0$ | 20.41 | 32.1 | 0.483 | -6.487 | 14.902 | 0.429 | -0.88 | 43.98 | 0.435 | -7.365 | 29.788 | 0.374 |
| $f_{NEA} = 0;\ f_{DEN} = 0.005$ | 29.13 | 5.53 | 0.465 | -4.662 | 1.019 | 0.48 | 78.32 | 60.3 | 0.473 | 65.714 | 69.991 | 0.486 |
| $f_{NEA} = 0;\ f_{DEN} = 0.01$ | -23.97 | 21.18 | 0.433 | 5.839 | 1.002 | 0.484 | 113.22 | 102.37 | 0.484 | 141.515 | 115.508 | 0.411 |
| $f_{NEA} = 0;\ f_{DEN} = 0.015$ | -6.94 | -20.55 | 0.479 | 13.673 | 9.471 | 0.484 | 219.63 | 186.55 | 0.453 | 216.295 | 211.303 | 0.481 |
| $f_{NEA} = 0;\ f_{DEN} = 0.02$ | 2.11 | -9.7 | 0.48 | 2.425 | 21.277 | 0.446 | 283.82 | 264.25 | 0.472 | 282.137 | 288.326 | 0.482 |
| $f_{NEA} = 0.005;\ f_{DEN} = 0$ | 139.01 | 163.2 | 0.464 | 154.998 | 168.042 | 0.453 | 358.62 | 358.98 | 0.499 | 373.579 | 389.946 | 0.442 |
| $f_{NEA} = 0.005;\ f_{DEN} = 0.005$ | 143.59 | 149.68 | 0.491 | 133.738 | 118.959 | 0.45 | 440.15 | 462.33 | 0.465 | 417.926 | 404.327 | 0.454 |
| $f_{NEA} = 0.005;\ f_{DEN} = 0.01$ | 129.2 | 100.53 | 0.463 | 134.032 | 144.062 | 0.462 | 509.87 | 476.26 | 0.456 | 486.115 | 494.417 | 0.472 |
| $f_{NEA} = 0.005;\ f_{DEN} = 0.015$ | 140.21 | 109.42 | 0.456 | 149.708 | 151.293 | 0.494 | 584.32 | 604.06 | 0.471 | 568.648 | 564.759 | 0.488 |
| $f_{NEA} = 0.005;\ f_{DEN} = 0.02$ | 149.03 | 171.9 | 0.464 | 140.381 | 140.525 | 0.5 | 649.73 | 659.92 | 0.487 | 619.205 | 622.525 | 0.489 |
| $f_{NEA} = 0.01;\ f_{DEN} = 0$ | 305.63 | 339.3 | 0.456 | 295.404 | 292.712 | 0.491 | 765.68 | 779.31 | 0.483 | 731.528 | 729.959 | 0.495 |
| $f_{NEA} = 0.01;\ f_{DEN} = 0.005$ | 276.31 | 269.7 | 0.49 | 295.794 | 291.864 | 0.486 | 790.26 | 767.4 | 0.467 | 796.144 | 789.836 | 0.477 |
| $f_{NEA} = 0.01;\ f_{DEN} = 0.01$ | 260.29 | 248.33 | 0.479 | 299.015 | 314.712 | 0.447 | 857.58 | 848.71 | 0.488 | 868.268 | 883.897 | 0.449 |
| $f_{NEA} = 0.01;\ f_{DEN} = 0.015$ | 297.87 | 300.06 | 0.496 | 290.962 | 284.05 | 0.474 | 930.87 | 934.52 | 0.494 | 926.772 | 915.821 | 0.459 |
| $f_{NEA} = 0.01;\ f_{DEN} = 0.02$ | 300.64 | 318.05 | 0.474 | 287.159 | 298.565 | 0.458 | 1005.36 | 975.48 | 0.459 | 1004.375 | 1038.56 | 0.391 |
| $f_{NEA} = 0.015;\ f_{DEN} = 0$ | 418.29 | 451.18 | 0.451 | 460.899 | 463.85 | 0.49 | 1144.69 | 1170.89 | 0.464 | 1116.882 | 1117.64 | 0.498 |
| $f_{NEA} = 0.015;\ f_{DEN} = 0.005$ | 408.56 | 428.37 | 0.47 | 432.525 | 439.239 | 0.478 | 1141.79 | 1131.85 | 0.485 | 1147.234 | 1143.431 | 0.488 |
| $f_{NEA} = 0.015;\ f_{DEN} = 0.01$ | 434.95 | 401.08 | 0.454 | 430.956 | 441.178 | 0.466 | 1184.71 | 1173.34 | 0.483 | 1198.455 | 1208.898 | 0.463 |
| $f_{NEA} = 0.015;\ f_{DEN} = 0.015$ | 493.5 | 481.98 | 0.485 | 419.808 | 423.074 | 0.489 | 1322.54 | 1283.41 | 0.44 | 1260.104 | 1262.36 | 0.493 |
| $f_{NEA} = 0.015;\ f_{DEN} = 0.02$ | 435.12 | 455.33 | 0.469 | 444.705 | 447.39 | 0.49 | 1368.13 | 1338.17 | 0.452 | 1352.822 | 1343.623 | 0.467 |
| $f_{NEA} = 0.02;\ f_{DEN} = 0$ | 571.4 | 566.62 | 0.492 | 585.948 | 570.366 | 0.446 | 1429.72 | 1449.12 | 0.47 | 1471.65 | 1467.111 | 0.483 |
| $f_{NEA} = 0.02;\ f_{DEN} = 0.005$ | 589.33 | 545.78 | 0.435 | 560.738 | 577.273 | 0.443 | 1561.06 | 1534.65 | 0.458 | 1503.25 | 1515.064 | 0.461 |
| $f_{NEA} = 0.02;\ f_{DEN} = 0.01$ | 599.1 | 561.22 | 0.443 | 557.286 | 563.83 | 0.477 | 1588.19 | 1547.05 | 0.439 | 1560.229 | 1564.305 | 0.486 |
| $f_{NEA} = 0.02;\ f_{DEN} = 0.015$ | 596.92 | 613.3 | 0.475 | 570.059 | 549.534 | 0.428 | 1664.67 | 1645.85 | 0.47 | 1624.164 | 1619.687 | 0.484 |
| $f_{NEA} = 0.02;\ f_{DEN} = 0.02$ | 636.11 | 622.51 | 0.479 | 608.509 | 590.207 | 0.435 | 1739.12 | 1768.77 | 0.452 | 1742.161 | 1730.27 | 0.46 |

Table S20: Assessing if *(ABBA − BABA) − (BAAA − ABAA)* significantly differs from zero in the hypothetical European and Asian populations under a multi-pulse model of introgression from Neanderthals and Denisovans with an additional pulse of gene flow from a so called "Basal Eurasian" population (see Figure S5) for all pairwise combinations of admixture proportions ($f_{NEA}$ and $f_{DEN}$) from 0% to 2%. The rows correspond to the simulated admixture proportions, the columns represent the mean site pattern difference across 100 replicate simulations and the corresponding *p-value* to assess if *(ABBA − BABA) − (BAAA − ABAA)* significantly differs from zero. After accounting for multiple-comparisons using the Bonferroni correction, a *p-value* less than 0.002 is considered significant.

| | $ABBA - BABA$ | $BAAA - ABAA$ | $p - value$ |
|---|---|---|---|
| CEU | 13960.242 | 14314.081 | 0.402 |
| FIN | 14240.051 | 16109.465 | 0.14 |
| GBR | 13621.099 | 14726.33 | 0.277 |
| IBS | 13375.0 | 15011.692 | 0.202 |
| TSI | 13279.486 | 14338.346 | 0.256 |
| CHB | 16789.359 | 19883.155 | 0.101 |
| CHS | 16618.8 | 20871.371 | 0.059 |
| CDX | 16470.989 | 20108.935 | 0.023 |
| JPT | 16341.356 | 19786.279 | 0.054 |
| KHV | 16281.626 | 19061.303 | 0.07 |
| BEB | 14977.07 | 18427.163 | 0.039 |
| GIH | 14534.204 | 17736.437 | 0.064 |
| ITU | 14647.108 | 17299.304 | 0.05 |
| PJL | 14412.208 | 16719.917 | 0.057 |
| STU | 14312.333 | 17161.294 | 0.053 |
| CLM | 13233.394 | 14108.734 | 0.301 |
| MXL | 13875.812 | 15756.422 | 0.145 |
| PEL | 14777.082 | 16675.824 | 0.125 |
| PUR | 11820.096 | 12472.481 | 0.335 |

Table S21: Assessing if *(ABBA − BABA) − (BAAA − ABAA)* significantly differs from zero for *((YRI, P2), Altai Neanderthal)* trios in the 1000 genomes project. Rows represent the *P2* population for the trio results and the columns the columns represent the mean site pattern difference across trios and the corresponding *p-value* to assess if *(ABBA − BABA) − (BAAA − ABAA)* significantly differs from zero. After accounting for multiple-comparisons using the Bonferroni correction, a *p-value* less than 0.0026 is considered significant.

| | $D$ | | $D_{anc}$ | | $D+$ | | $f_{hom}$ | | $f_{anc}$ | | $f+$ | |
|---|---|---|---|---|---|---|---|---|---|---|---|---|
| | $\mu$ | $\sigma$ | $\mu$ | $\sigma$ | $\mu$ | $\sigma$ | $\mu$ | $\sigma$ | $\mu$ | $\sigma$ | $\mu$ | $\sigma$ |
| CEU | 0.041 | 0.004 | 0.01 | 0.001 | 0.016 | 0.002 | 0.012 | 0.001 | 0.012 | 0.002 | 0.012 | 0.001 |
| FIN | 0.041 | 0.003 | 0.011 | 0.001 | 0.017 | 0.002 | 0.012 | 0.001 | 0.013 | 0.002 | 0.012 | 0.001 |
| GBR | 0.04 | 0.004 | 0.01 | 0.001 | 0.016 | 0.002 | 0.011 | 0.001 | 0.012 | 0.002 | 0.012 | 0.001 |
| IBS | 0.039 | 0.003 | 0.011 | 0.002 | 0.016 | 0.002 | 0.011 | 0.001 | 0.012 | 0.002 | 0.012 | 0.001 |
| TSI | 0.039 | 0.004 | 0.01 | 0.002 | 0.016 | 0.002 | 0.011 | 0.001 | 0.012 | 0.002 | 0.011 | 0.001 |
| CHB | 0.049 | 0.004 | 0.014 | 0.002 | 0.021 | 0.002 | 0.014 | 0.001 | 0.016 | 0.002 | 0.015 | 0.001 |
| CHS | 0.048 | 0.004 | 0.015 | 0.002 | 0.021 | 0.002 | 0.014 | 0.001 | 0.017 | 0.002 | 0.015 | 0.002 |
| CDX | 0.048 | 0.003 | 0.014 | 0.001 | 0.021 | 0.002 | 0.014 | 0.001 | 0.016 | 0.002 | 0.015 | 0.001 |
| JPT | 0.047 | 0.003 | 0.014 | 0.002 | 0.021 | 0.002 | 0.014 | 0.001 | 0.016 | 0.002 | 0.015 | 0.001 |
| KHV | 0.047 | 0.003 | 0.014 | 0.001 | 0.02 | 0.001 | 0.014 | 0.001 | 0.015 | 0.002 | 0.015 | 0.001 |
| BEB | 0.044 | 0.003 | 0.013 | 0.002 | 0.019 | 0.002 | 0.013 | 0.001 | 0.015 | 0.002 | 0.014 | 0.001 |
| GIH | 0.042 | 0.003 | 0.013 | 0.002 | 0.019 | 0.002 | 0.012 | 0.001 | 0.014 | 0.002 | 0.013 | 0.001 |
| ITU | 0.043 | 0.003 | 0.012 | 0.001 | 0.018 | 0.001 | 0.012 | 0.001 | 0.014 | 0.001 | 0.013 | 0.001 |
| PJL | 0.042 | 0.003 | 0.012 | 0.001 | 0.018 | 0.002 | 0.012 | 0.001 | 0.013 | 0.002 | 0.013 | 0.001 |
| STU | 0.042 | 0.004 | 0.012 | 0.002 | 0.018 | 0.002 | 0.012 | 0.001 | 0.014 | 0.002 | 0.013 | 0.001 |
| CLM | 0.039 | 0.005 | 0.01 | 0.002 | 0.016 | 0.002 | 0.011 | 0.001 | 0.011 | 0.002 | 0.011 | 0.001 |
| MXL | 0.04 | 0.004 | 0.011 | 0.001 | 0.017 | 0.002 | 0.012 | 0.001 | 0.013 | 0.002 | 0.012 | 0.001 |
| PEL | 0.043 | 0.004 | 0.012 | 0.002 | 0.018 | 0.002 | 0.012 | 0.001 | 0.013 | 0.002 | 0.013 | 0.001 |
| PUR | 0.035 | 0.005 | 0.009 | 0.002 | 0.014 | 0.002 | 0.01 | 0.001 | 0.01 | 0.002 | 0.01 | 0.002 |

Table S22: The mean ($\mu$) and standard deviation ($\sigma$) of all introgression metrics for *((YRI, P2), Altai Neanderthal)* trios in the 1000 genomes project where rows represents the respective *P2* population for the trio results.

| | Site Pattern Counts | | | | Site Pattern Differences | | Detection Metrics | | | | | | Quantification Metrics | | | | | |
|---|---|---|---|---|---|---|---|---|---|---|---|---|---|---|---|---|---|---|
| | $ABBA$ | $BABA$ | $BAAA$ | $ABAA$ | $ABBA - BABA$ | $BAAA - ABAA$ | $D$ | $p - value$ | $D_{anc}$ | $p - value$ | $D+$ | $p - value$ | $f_{hom}$ | $\sigma$ | $f_{anc}$ | $\sigma$ | $f+$ | $\sigma$ |
| CEU | 178686.455 | 164683.266 | 710638.497 | 695172.877 | 14003.189 | 15465.62 | 0.041 | 0.0 | 0.011 | 0.0 | 0.017 | 0.0 | 0.013 | 0.002 | 0.013 | 0.002 | 0.013 | 0.002 |
| FIN | 179031.036 | 164668.561 | 710965.968 | 693572.578 | 14362.474 | 17393.39 | 0.042 | 0.0 | 0.012 | 0.0 | 0.018 | 0.0 | 0.013 | 0.002 | 0.015 | 0.002 | 0.014 | 0.001 |
| GBR | 178557.14 | 164879.991 | 710566.718 | 694572.368 | 13677.149 | 15994.35 | 0.04 | 0.0 | 0.011 | 0.0 | 0.017 | 0.0 | 0.012 | 0.002 | 0.014 | 0.002 | 0.013 | 0.002 |
| IBS | 178070.734 | 164607.78 | 709977.626 | 693780.074 | 13462.954 | 16197.552 | 0.039 | 0.0 | 0.012 | 0.0 | 0.017 | 0.0 | 0.012 | 0.002 | 0.014 | 0.002 | 0.013 | 0.001 |
| TSI | 178060.36 | 164743.257 | 710026.979 | 694487.596 | 13317.104 | 15539.384 | 0.039 | 0.0 | 0.011 | 0.0 | 0.017 | 0.0 | 0.012 | 0.002 | 0.013 | 0.002 | 0.013 | 0.002 |
| CHB | 181002.098 | 164195.934 | 712256.723 | 691202.309 | 16806.164 | 21054.414 | 0.049 | 0.0 | 0.015 | 0.0 | 0.022 | 0.0 | 0.015 | 0.002 | 0.018 | 0.002 | 0.017 | 0.002 |
| CHS | 180875.077 | 164219.927 | 712291.95 | 690251.844 | 16655.15 | 22040.106 | 0.048 | 0.0 | 0.016 | 0.0 | 0.022 | 0.0 | 0.015 | 0.002 | 0.019 | 0.002 | 0.017 | 0.002 |
| CDX | 180749.697 | 164233.596 | 712372.16 | 691096.065 | 16516.101 | 21276.095 | 0.048 | 0.0 | 0.015 | 0.0 | 0.022 | 0.0 | 0.015 | 0.002 | 0.018 | 0.002 | 0.017 | 0.002 |
| JPT | 180807.724 | 164447.848 | 712199.429 | 691255.526 | 16359.876 | 20943.903 | 0.047 | 0.0 | 0.015 | 0.0 | 0.021 | 0.0 | 0.015 | 0.002 | 0.018 | 0.002 | 0.016 | 0.002 |
| KHV | 180684.322 | 164336.497 | 712244.409 | 692051.098 | 16347.825 | 20193.311 | 0.047 | 0.0 | 0.014 | 0.0 | 0.021 | 0.0 | 0.015 | 0.002 | 0.017 | 0.002 | 0.016 | 0.002 |
| BEB | 179409.557 | 164398.409 | 711619.425 | 692038.09 | 15011.148 | 19581.335 | 0.044 | 0.0 | 0.014 | 0.0 | 0.02 | 0.0 | 0.013 | 0.001 | 0.017 | 0.002 | 0.015 | 0.001 |
| GIH | 178956.411 | 164389.555 | 711067.865 | 692115.643 | 14566.856 | 18952.223 | 0.042 | 0.0 | 0.014 | 0.0 | 0.019 | 0.0 | 0.013 | 0.001 | 0.016 | 0.002 | 0.015 | 0.001 |
| ITU | 179084.768 | 164362.731 | 711261.704 | 692726.341 | 14722.036 | 18535.362 | 0.043 | 0.0 | 0.013 | 0.0 | 0.019 | 0.0 | 0.013 | 0.001 | 0.016 | 0.001 | 0.015 | 0.001 |
| PJL | 178926.57 | 164445.326 | 711200.889 | 693279.051 | 14481.244 | 17921.838 | 0.042 | 0.0 | 0.013 | 0.0 | 0.019 | 0.0 | 0.013 | 0.001 | 0.015 | 0.001 | 0.014 | 0.001 |
| STU | 178819.539 | 164423.601 | 710873.484 | 692550.773 | 14395.938 | 18322.711 | 0.042 | 0.0 | 0.013 | 0.0 | 0.019 | 0.0 | 0.013 | 0.001 | 0.016 | 0.001 | 0.014 | 0.001 |
| CLM | 177906.744 | 164638.873 | 709986.837 | 694691.111 | 13267.871 | 15295.726 | 0.039 | 0.0 | 0.011 | 0.0 | 0.016 | 0.0 | 0.012 | 0.001 | 0.013 | 0.001 | 0.013 | 0.001 |
| MXL | 178805.656 | 164842.313 | 710971.409 | 694089.274 | 13963.343 | 16882.135 | 0.041 | 0.0 | 0.012 | 0.0 | 0.018 | 0.0 | 0.013 | 0.001 | 0.015 | 0.001 | 0.014 | 0.001 |
| PEL | 179596.91 | 164775.627 | 711624.143 | 693882.313 | 14821.283 | 17741.831 | 0.043 | 0.0 | 0.013 | 0.0 | 0.019 | 0.0 | 0.013 | 0.002 | 0.015 | 0.002 | 0.014 | 0.002 |
| PUR | 176552.434 | 164690.52 | 708663.714 | 694972.186 | 11861.914 | 13691.528 | 0.035 | 0.0 | 0.01 | 0.0 | 0.015 | 0.0 | 0.011 | 0.001 | 0.012 | 0.001 | 0.011 | 0.001 |

Table S23: All site pattern counts and estimates of introgression metrics calculated from derived allele frequencies among non-African populations in the 1000 Genome's Project. All *p-values* and standard deviations ($\sigma$) were calculated from a bootstrapped distribution consisting of 1000 bootstrapped replicates. After accounting for multiple-comparisons using the Bonferroni correction, a *p-value* less than 0.0026 is considered significant.

| | Site Pattern Counts | | | | Site Pattern Differences | | Detection Metrics | | | | | | Quantification Metrics | | | | | |
|---|---|---|---|---|---|---|---|---|---|---|---|---|---|---|---|---|---|---|
| | $ABBA$ | $BABA$ | $BAAA$ | $ABAA$ | $ABBA - BABA$ | $BAAA - ABAA$ | $D$ | $p - value$ | $D_{anc}$ | $p - value$ | $D+$ | $p - value$ | $f_{hom}$ | $\sigma$ | $f_{anc}$ | $\sigma$ | $f+$ | $\sigma$ |
| (((Dingo, Basenji), Israeli Wolf), Golden Jackal) | 186412.0 | 154645.0 | 413597.0 | 370899.0 | 31767.0 | 42698.0 | 0.093 | 0.0 | 0.054 | 0.0 | 0.066 | 0.0 | 0.051 | 0.004 | 0.065 | 0.005 | 0.058 | 0.004 |

Table S24: All site pattern counts and estimates of introgression metrics from single whole-genome sequences for the trio ((Dingo, Basenji), Israeli Wolf) where the Golden Jackal was used to polarize ancestral states. All *p-values* and standard deviations ($\sigma$) were calculated from a bootstrapped distribution consisting of 1000 bootstrapped replicates. A *p-value* less than 0.05 is considered significant.
